# Disorder is a critical component of lipoprotein sorting in Gram-negative bacteria

**DOI:** 10.1101/2021.01.05.425367

**Authors:** Jessica El Rayes, Joanna Szewczyk, Michael Deghelt, André Matagne, Bogdan I. Iorga, Seung-Hyun Cho, Jean-François Collet

**Author notes:** Both authors contributed equally to the work (Jessica El Rayes, Joanna Szewczyk).

## Abstract

Gram-negative bacteria express structurally diverse lipoproteins in their envelope. Here we found that approximately half of lipoproteins destined to the *Escherichia coli* outer membrane display an intrinsically disordered linker at their N-terminus. Intrinsically disordered regions are common in proteins, but establishing their importance *in vivo* has remained challenging. Here, as we sought to unravel how lipoproteins mature, we discovered that unstructured linkers are required for optimal trafficking by the Lol lipoprotein sorting system: linker deletion re-routes three unrelated lipoproteins to the inner membrane. Focusing on the stress sensor RcsF, we found that replacing the linker with an artificial peptide restored normal outer membrane targeting only when the peptide was of similar length and disordered. Overall, this study reveals the role played by intrinsic disorder in lipoprotein sorting, providing mechanistic insight into the biogenesis of these proteins and suggesting that evolution can select for intrinsic disorder that supports protein function.

## Introduction

The cell envelope is the morphological hallmark of *Escherichia coli* and other Gram-negative bacteria. It is composed of the inner membrane, a classical phospholipid bilayer, as well as the outer membrane, an asymmetric bilayer with phospholipids in the inner leaflet and lipopolysaccharides in the outer leaflet^1^. This lipid asymmetry enables the outer membrane to function as a barrier that effectively prevents the diffusion of toxic compounds in the environment into the cell. The inner and outer membranes are separated by the periplasm, a viscous compartment that contains a thin layer of peptidoglycan also known as the cell wall^1^. The cell envelope is essential for growth and survival, as illustrated by the fact that several antibiotics such as the β-lactams target mechanisms of envelope assembly. Mechanisms involved in envelope biogenesis and maintenance are therefore attractive targets for novel antibacterial strategies.

Approximately one-third of *E. coli* proteins are targeted to the envelope, either as soluble proteins present in the periplasm or as proteins inserted in one of the two membranes^2^. While inner membrane proteins cross the lipid bilayer via one or more hydrophobic α-helices, proteins inserted in the outer membrane generally adopt a β-barrel conformation^3^. Another important group of envelope proteins is the lipoproteins, which are globular proteins anchored to one of the two membranes by a lipid moiety. Lipoproteins carry out a variety of important functions in the cell envelope: they participate in the biogenesis of the outer membrane by inserting lipopolysaccharide molecules^4,5^ and β-barrel proteins^6^, they function as stress sensors triggering signal transduction cascades when envelope integrity is altered^7^, and they control processes that are important for virulence^8^. The diverse roles played by lipoproteins in the cell envelope has drawn a lot of attention lately, revealing how crucial these proteins are in a wide range of vital processes and identifying them as attractive targets for antibiotic development. Yet, a detailed understanding of the mechanisms involved in lipoprotein maturation and trafficking is still missing.

Lipoproteins are synthesized in the cytoplasm as precursors with an N-terminal signal peptide^9^. The last four C-terminal residues of this signal peptide, known as the lipobox, function as a molecular determinant of lipid modification unique to bacteria; only the cysteine at the last position of the lipobox is strictly conserved^10^. After secretion of the lipoprotein into the periplasm, the thiol side-chain of the cysteine is first modified with a diacylglyceryl moiety by prolipoprotein diacylglyceryl transferase (Lgt)^9^ (**Extended Data Fig. 1a**, step 1). Then, signal peptidase II (LspA) catalyzes cleavage of the signal peptide N-terminally of the lipidated cysteine before apolipoprotein N-acyltransferase (Lnt) adds a third acyl group to the N-terminal amino group of the cysteine (**Extended Data Fig. 1a**, steps 2-3). Most mature lipoproteins are then transported to the outer membrane by the Lol system. Lol consists of LolCDE, an ABC transporter that extracts lipoproteins from the inner membrane and transfers them to the soluble periplasmic chaperone LolA (**Extended Data Fig. 1a**, steps 4-5)^11^. LolA escorts lipoproteins across the periplasm, binding their hydrophobic lipid tail, and delivers them to the outer membrane lipoprotein LolB (**Extended Data Fig. 1a**, step 6). LolB finally anchors lipoproteins to the inner leaflet of the outer membrane using a mechanism that remains poorly characterized (**Extended Data Fig. 1a**, step 7).

In most Gram-negative bacteria, a few lipoproteins remain in the inner membrane^12,13^. The current view is that inner membrane retention depends on the identity of the two residues located immediately downstream of the N-terminal cysteine on which the lipid moiety is attached^14^; this sequence, two amino acids in length, is known as the Lol sorting signal. When lipoproteins have an aspartate at position +2 and an aspartate, glutamate, or glutamine at position +3, they remain in the inner membrane^15,16^, possibly because strong electrostatic interactions between the +2 aspartate and membrane phospholipids prevent their interaction with LolCDE^17^. However, this model is largely based on data obtained in *E. coli* and variations have been described in other bacteria. For instance, in the pathogen *Pseudomonas aeruginosa*, an aspartate is rarely found at position +2 and inner membrane retention appears to be determined by residues +3 and +4^18,19^. Surprisingly, lipoproteins are well sorted in *P. aeruginosa* cells expressing the *E. coli* LolCDE complex^20^, despite their different Lol sorting signal. This result cannot be explained by the current model of lipoprotein sorting, underscoring that our comprehension of the precise mechanism that governs the triage of lipoproteins remains incomplete.

Excitingly, more unresolved questions regarding lipoprotein biogenesis have recently been raised. First, it was reported that a LolA-LolB-independent trafficking route to the outer membrane exists in *E. coli*^21^, but the factors involved have remained unknown. Second, although lipoproteins have traditionally been considered to be exposed to the periplasm in *E. coli* and many other bacterial models^9^, a series of investigations have started to challenge this view by identifying lipoproteins on the surface of *E. coli, Vibrio cholerae*, and *Salmonella* Typhimurium^22–26^. Overall, the field is beginning to explore a lipoprotein topological landscape that is more complex than previously assumed and raising intriguing questions about the signals that control surface targeting and exposure.

Here, stimulated by the hypothesis that crucial details of the mechanisms underlying lipoprotein maturation remained to be elucidated, we sought to identify novel molecular determinants controlling lipoprotein biogenesis. First, we systematically analyzed the sequence of the 66 lipoproteins with validated localization^27^ encoded by the *E. coli* K12 genome^27^ and found that half of the outer membrane lipoproteins display a long and intrinsically disordered linker at their N-terminus. Intrigued by these unstructured segments, we then probed their importance for the biogenesis of RcsF, NlpD, and Pal, three structurally and functionally unrelated outer membrane lipoproteins. Unexpectedly, we found that deleting the linker—while keeping the Lol sorting signal intact—altered the targeting of all three lipoproteins to the outer membrane, with physiological consequences. Focusing on RcsF, we determined that both the length and disordered character of the linker were important. Remarkably, lowering the load of the Lol system by deleting *lpp*, which encodes the most abundant lipoprotein (~1 million copies per cell^28^), restored normal outer membrane targeting of linker-less RcsF, indicating that the N-terminal linker is required for optimal lipoprotein processing by Lol. Taken together, these observations reveal the unsuspected role played by protein intrinsic disorder in lipoprotein biogenesis.

## Results

### Half of *E. coli* lipoproteins present long disordered segments at their N-termini

In an attempt to discover novel molecular determinants controlling the biogenesis of lipoproteins, we decided to systematically analyze the sequence of the lipoproteins encoded by the *E. coli* genome (strain MG1655) in search of unidentified structural features. *E. coli* encodes ~80 validated lipoproteins^29^, of which 58 have been experimentally shown to localize in the outer membrane^27^. Comparative modeling of existing X-ray, cryogenic electron microscopy (cryo-EM), and nuclear magnetic resonance (NMR) structures revealed that approximately half of these outer membrane lipoproteins display a long segment (>22 residues) that is predicted to be disordered at the N-terminus (**Fig. 1, Extended Data Fig. 2, Extended Data Table 1**). In contrast, only one of the 8 lipoproteins that remain in the inner membrane (DcrB; **Extended Data Fig. 2, Extended Data Table 1**) had a long, disordered linker, suggesting that disordered peptides may be important for lipoprotein sorting.

**Figure 1.**
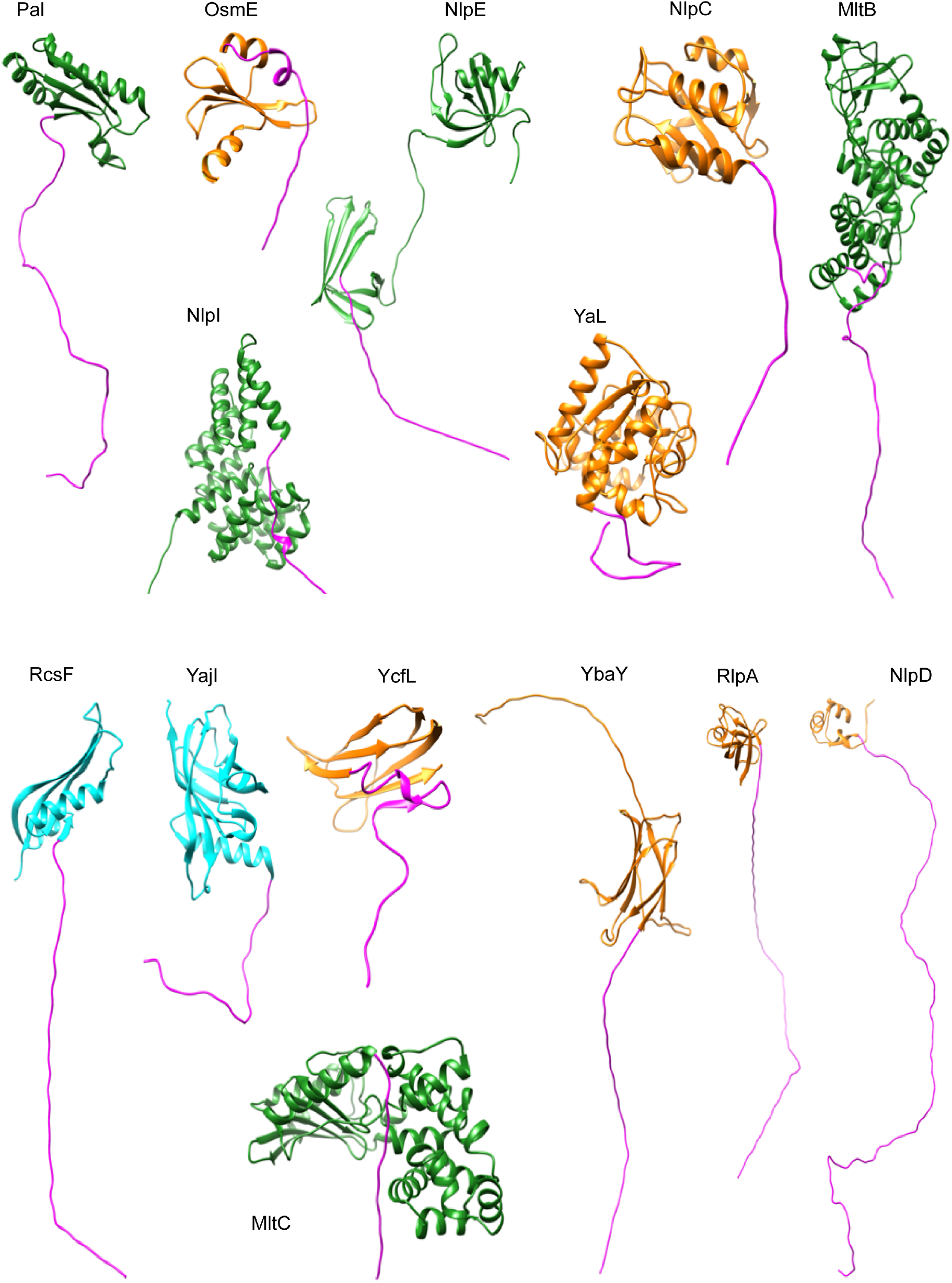
Structural analysis of lipoproteins reveals that half of outer membrane lipoproteins display an intrinsically disordered linker at the N-terminus. Structures were generated via comparative modeling (Methods). X-ray and cryo-EM structures are green, NMR structures are cyan, and structures built via comparative modeling from the closest analog in the same PFAM group are orange. In all cases, the N-terminal linker is magenta. Lipoproteins targeting the outer membrane: Pal, OsmE, NlpE, NlpC, MltB, NlpI, MltC, RcsF, YajI, YcfL, YbaY, RlpA, NlpD, YcaL. The 29 remaining lipoproteins are shown in Extended Data Figure 2.

### Deleting the N-terminal linker of RcsF, NlpD, and Pal perturbs their targeting to the outer membrane

Intrigued by the presence of these N-terminal disordered segments in so many outer membrane lipoproteins, we decided to investigate their functional importance. We selected three structurally unrelated lipoproteins whose function could easily be assessed: the stress sensor RcsF (which triggers the Rcs signaling cascade when damage occurs in the envelope^30^), NlpD (which activates the periplasmic N-acetylmuramyl-L-alanine amidase AmiC, which is involved in peptidoglycan cleavage during cell division^31,32^), and the peptidoglycan-binding lipoprotein Pal (which is important for outer membrane constriction during cell division^33^).

We began by preparing truncated versions of RcsF, NlpD, and Pal devoid of their N-terminal unstructured linkers (**Extended Data Fig. 1b, Extended Data Fig. 2;** RcsF_Δ19-47_, Pal_Δ26-56_, and NlpD_Δ29-64_). Note that the lipidated cysteine residue (+1) and the Lol sorting signal (the amino acids at positions +2 and +3) were not altered in RcsF_Δ19-47_, Pal_Δ26-56_, and NlpD_Δ29-64_, nor in any of the constructs discussed below (**Extended Data Table 2**). For Pal, although the unstructured linker spans residues 25-68 (**Fig. 1**), we used Pal_Δ26-56_ because Pal_Δ25-68_ was either degraded or not detected by the antibody (data not shown). We first tested whether the truncated lipoproteins were still correctly extracted from the inner membrane and transported to the outer membrane. The membrane fraction was prepared from cells expressing the three variants independently, and the outer and inner membranes were separated using sucrose density gradients (Methods). Whereas wild-type RcsF, NlpD, and Pal were mostly detected (>90%) in the outer membrane fraction, as expected, ~50% of RcsF_Δ19-47_ and ~60% of NlpD_Δ29-64_ were retained in the inner membrane (**Fig. 2a, 2b**). The sorting of Pal was also affected, although to a lesser extent: 15% of Pal_Δ26-56_ was retained in the inner membrane (**Fig. 2c**). Notably, the expression levels of the three linker-less variants were similar (NlpD_Δ29-64_) or lower (RcsF_Δ19-47_; Pal_Δ26-56_) than those of the wild-type proteins (**Extended Data Fig. 3**), indicating that accumulation in the inner membrane did not result from increased protein abundance.

**Figure 2.**
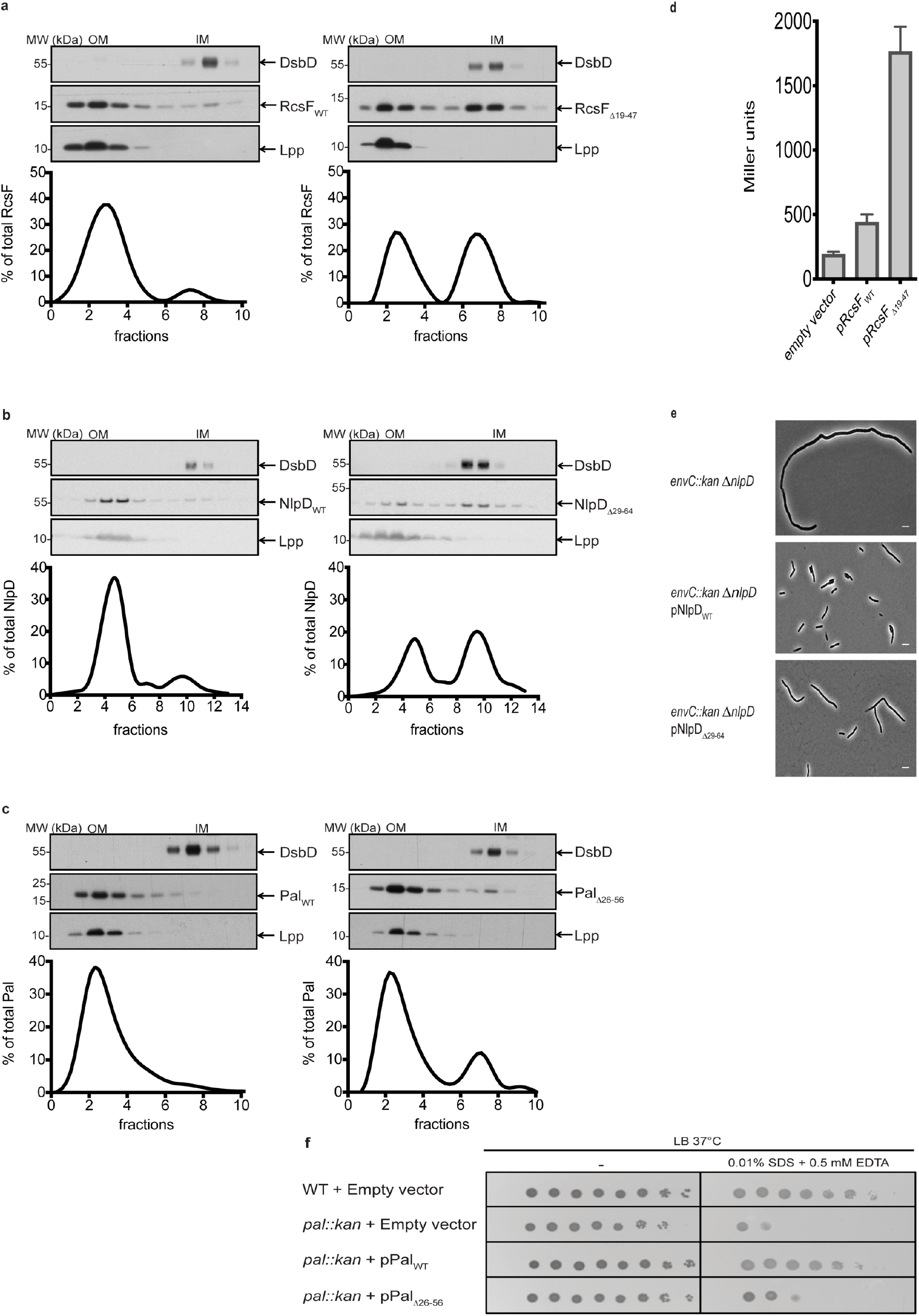
The N-terminal linker displayed by lipoproteins is important for outer membrane targeting. **a, b, c.** The outer membrane (OM) and inner membrane (IM) were separated via centrifugation in a three-step sucrose density gradient (Methods). While (c) RcsFWT, (d) NlpDWT, and (e) Pal_WT_ were found predominantly in the OM, RcsF_Δ19-47_, NlpD_Δ29-64_, and Pal_Δ26-56_ were substantially retained in the IM. Data are presented as the ratio of signal intensity in a single fraction to the total intensity in all fractions. All variants were expressed from plasmids (**Extended Data Table 4**). DsbD and Lpp were used as controls for the OM and IM, respectively. **d.** The Rcs system is constitutively active when RcsF’s linker is missing. Rcs activity was measured with a beta-galactosidase assay in a strain harboring a transcriptional *rprA::lacZ* fusion (Methods). Results were normalized to expression levels of RcsF variants (mean ± standard deviation; n = 6 biologically independent experiments) **e.** Phase-contrast images of the *envC::kan* Δ*nlpD* mutant complemented with NlpD_WT_ or NlpD_Δ29-64_. NlpD_Δ29-64_ only partially rescues the chaining phenotype of the *envC::kan* Δ*nlpD* double mutant. Scale bar, 5 μm. **f.** Expression of Pal_Δ26-56_ does not rescue the sensitivity of the *pal::kan* mutant to SDS-EDTA. Cells were grown in LB medium at 37 °C until OD_600_ = 0.5. Tenfold serial dilutions were made in LB, plated onto LB agar or LB agar supplemented with 0.01% SDS and 0.5 mM EDTA, and incubated at 37 °C. Images in **a**, **b**, **c**, **e**, and **f** are representative of biological triplicates. Graphs in **a**, **b**, and **c** were created by spline analysis of curves representing a mean of three independent experiments.

We then tested the impact of linker deletion on the function of these three proteins. In cells expressing RcsF_Δ19-47_, the Rcs system was constitutively turned on (**Fig. 2d**); when RcsF accumulates in the inner membrane, it becomes available for interaction with IgaA, its downstream Rcs partner in the inner membrane^30,34^. Likewise, expression of NlpDΔ29-64 did not rescue the chaining phenotype (**Fig. 2e**)^35^ exhibited by cells lacking both *nlpD* and *envC*, an activator of the amidases AmiA and AmiB^32^. Finally, Pal_Δ26-56_ partially rescued the sensitivity of the *pal* mutant to SDS-EDTA that results from increased membrane permeability^36^ (**Fig. 2f**). However, this observation needs to be considered with caution given that Pal_Δ26-56_ seemed to be expressed at lower levels than wild-type Pal (**Extended Data Fig. 3**). Thus, preventing normal targeting of RcsF, NlpD and Pal to the outer membrane had functional consequences.

### RcsF variants with unstructured artificial linkers of similar lengths are normally targeted to the outer membrane

The results above were surprising because they revealed that the normal targeting of RcsF, NlpD, and Pal to the outer membrane does not only require an appropriate Lol sorting signal, as proposed by the current model for lipoprotein sorting^9^, but also the presence of an N-terminal linker. We selected RcsF, whose accumulation in the inner membrane can be easily tracked by monitoring Rcs activity^30,37^, to investigate the structural features of the linker controlling lipoprotein maturation; keeping as little as 10% of the total pool of RcsF molecules in the inner membrane is sufficient to fully activate Rcs^30^.

We first tested whether changing the sequence of the N-terminal segment while preserving its disordered character still yielded normal targeting of the protein to the outer membrane. To that end, we prepared an RcsF variant in which the N-terminal linker was replaced by an artificial, unstructured sequence (**Extended Data Table 2, Extended Data Fig. 2, Extended Data Fig. 4**) of similar length and consisting mostly of GS repeats (RcsF_GS_). Substituting the wild-type linker with this artificial sequence was remarkably well tolerated by RcsF: RcsFGS was targeted normally to the outer membrane (**Fig. 3a**) and did not constitutively activate the stress system (**Fig. 3b**). Thus, although RcsFGS has an N-terminus with a completely different primary structure, it behaved like the wild-type protein.

**Figure 3.**
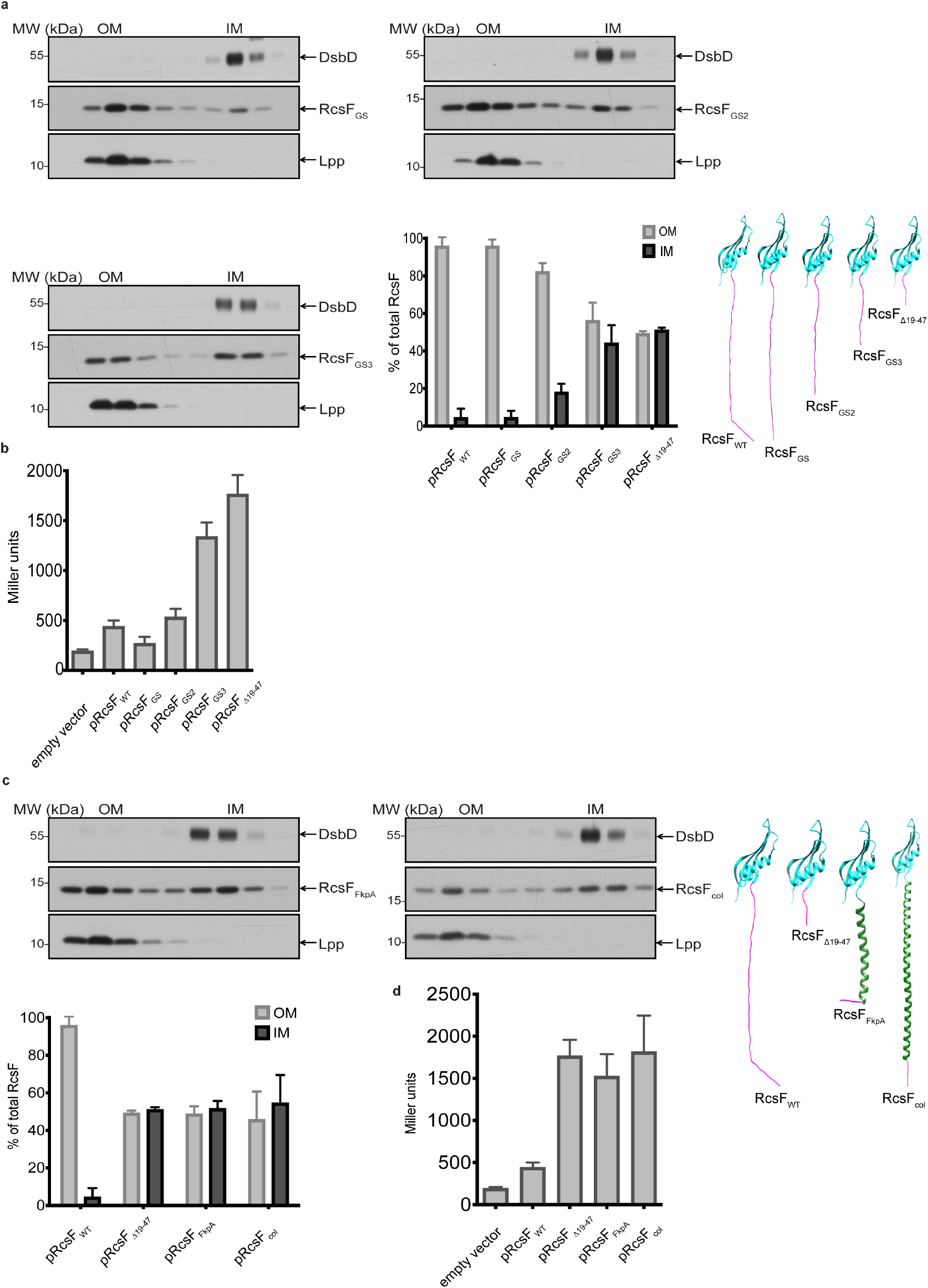
The length and the disordered character of the RcsF linker play key roles in RcsF targeting to the outer membrane. **a.** The outer membrane (OM) and inner membrane (IM) were separated via centrifugation in a three-step sucrose density gradient (Methods). DsbD and Lpp were used as controls for the OM and IM, respectively. The longer the linker, the more protein was correctly translocated to the IM. Bar graphs denote mean ± standard deviation of n = 3 biologically independent experiments. Images are representative of experiments and immunoblots performed in biological triplicate. **b.** Rcs activity was measured with a beta-galactosidase assay in a strain harboring a transcriptional *rprA::lacZ* fusion (Methods). Results were normalized to expression levels of RcsF variants (mean ± standard deviation of n = 6 biologically independent experiments). Rcs activity relates to the quantity of RcsF retained in the inner membrane. **c.** RcsF mutants harboring alpha helical linkers (RcsF_FkpA_ and RcsF_col_) were subjected to two consecutive centrifugations in sucrose density gradients (Methods). Both mutants were inefficiently translocated from the IM to the OM (mean ± standard deviation of n = 3 biologically independent experiments). Images are representative of experiments and immunoblots performed in biological triplicate. **d.** The Rcs system was constitutively active in RcsF_FkpA_ and RcsF_col_ strains; activation levels were comparable to those of RcsF_Δ19-47_. Rcs activity was measured as in **b**. Results were normalized as in **b**.

We then investigated whether the N-terminal linker required a minimal length for proper targeting and function. We therefore constructed two RcsF variants with shorter, unstructured, artificial linkers (RcsFGS2 and RcsFGS3, with linkers of 18 and 10 residues, respectively; **Extended Data Table 2, Extended Data Fig. 2, Extended Data Fig. 4**). Importantly, RcsFGS2 and, to a greater extent, RcsFGS3 did not properly localize to the outer membrane: the shorter the linker, the more RcsF remained in the inner membrane (**Fig. 3a**). Consistent with the amount of RcsFGS2 and RcsFGS3 retained in the inner membrane, Rcs activation levels were inversely related to linker length (**Fig. 3b**).

### The disordered character of the linker is required for normal targeting

Taken together, the results above demonstrated that the RcsF linker can be replaced with an artificial sequence lacking secondary structure, provided that it is of appropriate length. Next, we sought to directly probe the importance of having a disordered linker by replacing the RcsF linker with an alpha-helical segment 35 amino acids long from the periplasmic chaperone FkpA (RcsF_FkpA_; **Extended Data Table 2, Extended Data Fig. 2, Extended Data Fig. 4**). Introducing order at the N-terminus of RcsF dramatically impacted the protein distribution between the two membranes: RcsFFkpA was substantially retained in the inner membrane (**Fig. 3c**) and constitutively activated Rcs (**Fig. 3d**). As alpha-helical segments are considerably shorter than unstructured sequences containing a similar number of amino acids, we also prepared an RcsF variant (RcsF_col_) with a longer alpha helix from the helical segment of colicin Ia, which is 73 amino acids in length and also predicted to remain folded in the RcsF_col_ construct (**Extended Data Table 2, Extended Data Fig. 2, Extended Data Fig. 4**). However, doubling the size of the helix had no impact, with RcsFcol behaving similarly to RcsFFkpA (**Fig. 3c, 3d**). Together, these data demonstrate that having an N-terminal disordered linker downstream of the Lol sorting signal is required to correctly target RcsF to the outer membrane. The length of the linker is important, but the sequence is not, on the condition that the linker does not fold into a defined secondary structure.

### The disordered linker is required for optimal processing by Lol

Our finding that N-terminal disordered linkers function as molecular determinants of the targeting of lipoproteins to the outer membrane raised the question of whether these linkers work in a Lol-dependent or Lol-independent manner. To address this mechanistic question, we tested the impact of deleting *lpp* on the targeting of RcsF_Δ19-47_. The lipoprotein Lpp, also known as the Braun lipoprotein, covalently tethers the outer membrane to the peptidoglycan and controls the size of the periplasm ^38,39^. Being expressed at ~1 million copies per cell^28^, Lpp is numerically the most abundant protein in *E. coli*. Thus, by deleting *lpp*, we considerably decreased the load on the Lol system by removing its most abundant substrate. Remarkably, *lpp* deletion fully rescued the targeting of RcsF_Δ19-47_ to the outer membrane (**Fig. 4a**), indicating that the linker functions in a Lol-dependent manner and suggesting that accumulation of RcsF_Δ19-47_ in the inner membrane results from a decreased ability of the Lol system to process the linker-less RcsF variant. Importantly, similar results were obtained with NlpDΔ29-64, which was also correctly targeted to the outer membrane in cells lacking Lpp (**Fig. 4a**). PalΔ26-56 could not be tested because membrane fractionation failed with *lpp pal* double mutant cells whether or not they expressed Pal_Δ26-56_ (data not shown).

**Figure 4.**
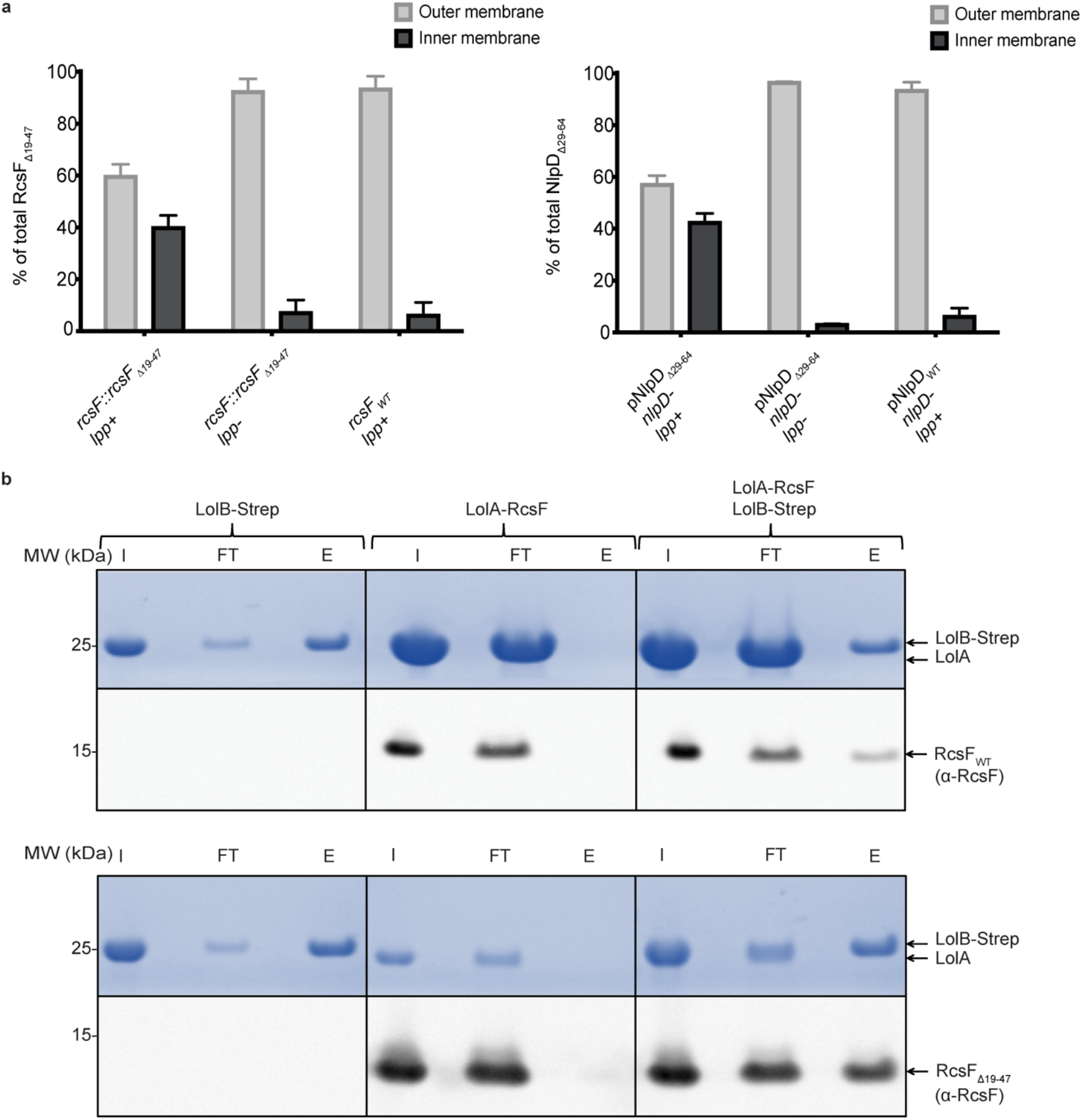
N-terminal disordered linkers interact with the Lol system to target lipoproteins to the outer membrane. **a.** Deleting Lpp rescues normal targeting of RcsF_Δ19-47_ and NlpD_Δ29-64_ to the outer membrane. The outer and inner membranes were separated via centrifugation in a sucrose density gradient (Methods). Whereas RcsF_Δ19-47_ and NlpD_Δ29-64_ accumulate in the inner membrane of cells expressing Lpp, the most abundant Lol substrate, they are normally targeted to the outer membrane in cells lacking Lpp (mean ± standard deviation of n = 3 biologically independent experiments). **b.** *In vitro* pull-down experiments show that RcsF_WT_ and RcsF_Δ19-47_ are transferred from LolA to LolB. LolA-RcsF_WT_ and LolA-RcsF_Δ19-47_ complexes were obtained by LolA-His affinity chromatography followed by size exclusion chromatography (Methods). Each complex was incubated with LolB-Strep that was previously purified via Strep-Tactin affinity chromatography (Methods). Both RcsF variants were eluted in complex with LolB-strep, while LolA was only present in the flow through. I, input; FT, flow through; E, eluate.

To obtain further insights into the mechanism at play here, we next monitored whether linker deletion impacted the transfer of RcsF from LolA to LolB *in vitro*. LolA with a C-terminal His-tag was expressed in the periplasm of cells expressing wild-type RcsF or RcsF_Δ19-47_ and purified to near homogeneity via affinity chromatography (Methods; **Extended Data Fig. 5**). Both RcsF and RcsF_Δ19-47_ were detected in immunoblots of the fractions containing purified LolA (**Extended Data Fig. 5**), indicating that both proteins form a soluble complex with LolA and confirming that they use this chaperone for transport across the periplasm. LolB was expressed as a soluble protein in the cytoplasm and purified by taking advantage of a C-terminal Strep-tag; LolB was then incubated with LolA-RcsF or LolA-RcsF_Δ19_._47_ and pulled-down using Streptactin beads (Methods). As both RcsF and RcsF_Δ19_._47_ were detected in the LolB-containing pulled-down fractions (**Fig. 4b**), we conclude that both proteins were transferred from LolA to LolB. Thus, the linker is not required for the transfer of RcsF from LolA to LolB.

Finally, we focused on the LolCDE ABC transporter in charge of extracting outer membrane lipoproteins and transferring them to LolA. Over-expression (**Extended Data Fig. 6a**) of all components of this complex failed to rescue normal targeting of RcsF_Δ19-47_ to the outer membrane (**Extended Data Fig. 6b**). Likewise, over-expressing the enzymes involved in lipoprotein maturation (Lgt, LspA, and Lnt; **Fig. 1**) had no impact on membrane targeting (**Extended Data Fig. 7a, 7b**). Thus, taken together, our results suggest that retention of RcsF_Δ19-47_ in the inner membrane does not result from the impairment of a specific step, but rather from less efficient processing of the truncated lipoprotein by the entire lipoprotein maturation pathway (see Discussion).

## Discussion

Lipoproteins are crucial for essential cellular processes such as envelope assembly and virulence. However, despite their functional importance and their potential as targets for new antibacterial therapies, we only have a vague understanding of the molecular factors that control their biogenesis. By discovering the role played by N-terminal disordered linkers in lipoprotein sorting, this study adds an important new layer to our comprehension of lipoprotein biogenesis in Gram-negative bacteria. Critically, it also indicates that the current model of lipoprotein sorting—that sorting between the two membranes is controlled by the 2 or 3 residues that are adjacent to the lipidated cysteine^40^—needs to be revised. Lipoproteins with unstructured linkers at their N-terminus are commonly found in Gram-negative bacteria including many pathogens (see below); further work will be required to determine whether these linkers control lipoprotein targeting in organisms other than *E. coli*, laying the foundation for designing new antibiotics.

It was previously shown that both *lolA* and *lolB* (but not *lolCDE*) can be deleted under specific conditions^21^, suggesting at least one alternate route for the transport of lipoproteins across the periplasm and their delivery to the outer membrane. During this investigation, we envisaged the possibility that the linker could be required to transport lipoproteins via a yet-to-be-identified pathway independent of LolA/LolB. However, our observations that both RcsF and RcsF_Δ19-47_ were found in complex with LolA **(Extended Data Fig. 5)** and were transferred by LolA to LolB **(Fig. 4b)** does not support this hypothesis. Instead, our data clearly indicate that lipoproteins with N-terminal linkers still depend on the Lol system for extraction from the inner membrane and transport to the outer membrane **(Extended Data Fig. 1a)**; they also suggest that N-terminal linkers improve lipoprotein processing by Lol (see below).

We note that two of the lipoproteins under investigation here, Pal and RcsF, have been reported to be surface-exposed^30,41,42^. A topology model has been proposed to explain how RcsF reaches the surface: the lipid moiety of RcsF is anchored in the outer leaflet of the outer membrane while the N-terminal linker is exposed on the cell surface before being threaded through the lumen of β-barrel proteins^42^. Thus, in this topology, the linker allows RcsF to cross the outer membrane. It is therefore tempting to speculate that N-terminal disordered linkers may be used by lipoproteins as a structural device to cross the outer membrane and reach the cell surface. It is worth noting that N-terminal linkers are commonly found in lipoproteins expressed by the pathogens *Borrelia burgdorferi* and *Neisseria meningitides*^24,43,44^; lipoprotein surface exposure is common in these pathogens. In addition, the accumulation of RcsF_Δ19-47_ in the inner membrane **(Fig. 2a)** also suggests that Lol may be using N-terminal linkers to recognize lipoproteins destined to the cell surface before their extraction from the inner membrane in order to optimize their targeting to the machinery exporting them to their final destination (BAM in the case of RcsF^30,42,45^). Investigating whether a dedicated Lol-dependent route exists for surface-exposed lipoproteins will be the subject of future research.

Our work also delivers crucial insights into the functional importance of disordered segments in proteins in general. Most proteins are thought to present portions that are intrinsically disordered. For instance, it is estimated that 30-50% of eukaryotic proteins contain regions that do not adopt a defined secondary structure *in vitro*^46^. However, demonstrating that these unstructured regions are functionally important *in vivo* is challenging. By showing that an N-terminal disordered segment downstream of the Lol signal is required for the correct sorting of lipoproteins, our work provides direct evidence that evolution has selected intrinsic disorder by function.

In conclusion, the data reported here establish that the triage of lipoproteins between the inner and outer membranes is not solely controlled by the Lol sorting signal; additional molecular determinants, such as protein intrinsic disorder, are also involved. Our data further highlight the previously unrecognized heterogeneity of the important lipoprotein family and call for a careful evaluation of the maturation pathways of these lipoproteins.

## DATA AVAILABILITY

All data generated or analysed during this study are included in this published article and its supplementary information file.

## ACKNOWLEDGMENTS

We thank Asma Boujtat for technical help. We are indebted to the members of the Collet laboratory and to Nassos Typas (EMBL, Heidelberg) for helpful suggestions and discussions and to Tom Silhavy (Princeton) for providing bacterial strains. J.S. was a research fellow of the FRIA and J.F.C. is an Investigator of the FRFS-WELBIO. This work was funded by the WELBIO, by grants from the F.R.S.-FNRS, from the Fédération Wallonie-Bruxelles (ARC 17/22-087), from the European Commission via the International Training Network Train2Target (721484), and from the EOS Excellence in Research Program of the FWO and FRS-FNRS (G0G0818N).

## AUTHOR CONTRIBUTIONS

J.-F.C., J.E.R., J.S., and S.H.C. designed and performed the experiments. J.E.R., J.S., and S.H.C. constructed the strains and cloned the constructs. J.-F.C., J.E.R., J.S., S.H.C., and A.M. analyzed and interpreted the data. B.I.I. performed the structural analysis. J.-F.C., J.E.R., and J.S. wrote the manuscript. All authors discussed the results and commented on the manuscript.

## METHODS

### Bacterial growth conditions

Bacterial strains used in this study are listed in **Extended Data Table 3**. Bacterial cells were cultured in Luria broth (LB) at 37 °C unless stated otherwise. The following antibiotics were added when appropriate: spectinomycin (100 μg/mL), ampicillin (200 μg/mL), chloramphenicol (25 μg/mL), and kanamycin (50 μg/mL). L-arabinose (0.2%) and isopropyl-β-D-thiogalactoside (IPTG) were used for induction when appropriate.

### Bacterial strains and plasmids

DH300 (a derivative of *Escherichia coli* MG1655 carrying a chromosomal *rprA::lacZ* fusion at the λ attachment site^47^) was used as wildtype throughout the study. All deletion mutants were obtained by transferring the corresponding alleles from the Keio collection^48^ (kan^R^) into DH300^47^ via P1 phage transduction. Deletions were verified by PCR and the absence of the protein was verified via immunoblotting (when possible). If necessary, the kanamycin cassette was removed via site-specific recombination mediated by the yeast Flp recombinase with pCP20 vector^49^. All strains expressing the RcsF mutants used for subcellular fractionation lacked *rcsB* in order to prevent induction of Rcs.

The plasmids used in this study are listed in **Extended Data Table 4** and the primers appear in **Extended Data Table 5**. RcsF, Pal, and NlpD were expressed from the low-copy vector pAM238^50^ containing the SC101 origin of replication and the *lac* promoter. To produce pSC202 for RcsF expression, *rcsF* (including approximately 30 base pairs upstream of the coding sequence) was amplified by PCR from the chromosome of DH300 (primer pair SH_RcsF(PstI)-R and SH_RcsFU-R (kpnI)-F). The amplification product was digested with KpnI and PstI and inserted into pAM238, resulting in pSC202. *nlpD* was amplified using primers JR1 and JR2 and *pal* was amplified with primers JS145 and JS146. Amplification products were digested with PstI-XbaI and KpnI-XbaI, respectively, generating pJR8 (for NlpD expression) and pJS20 (for Pal expression). To clone *rcsF_Δ19-47_*, the nucleotides encoding the RcsF signal sequence were amplified using primers SH_RcsFUR(kpnI)_F and SH_RcsFss-Fsg (NcoI)_R, and those encoding the RcsF signaling domain were amplified using primers SH_RcsFss-Fsg (NcoI)_R and SH_RcsF(PstI)_R. In both cases, pSC202 was used as template. Then, overlapping PCR was performed using SH_RcsFUR(kpnI)_F and SH_RcsF(PstI)_R from the two PCR products previously obtained. The final product was digested with KpnI and PstI, and ligated with pAM238 pre-digested with the same enzymes, yielding pSC201. To add a GS linker (Ser-Gly-Ser-Gly-Ser-Gly-Ala-Met) into pSC201, the primers SH_GS linker_F and SH_GS linker_R were mixed, boiled, annealed at room temperature, and ligated with pSC201 pre-digested with NcoI, generating pSC198. pSC199 was generated similarly, but using primers SH_SG linker_F and SH_SG linker_R and plasmid pSC198. pSC200 was generated using primers SH_Da linker_F and SH_SG linker_R and plasmid pSC199. The *pal* allele lacking the linker region (*pal_Δ26-56_*) was created via overlapping PCR. The pJS20 plasmid served as template for PCR with the M13R/M13F external primers and JS152/JS153 internal primers. The truncated allele was cloned into pAM238 at the same restriction sites as the full-length allele, producing pJS24. The *nlpD* allele lacking the linker regions (*nlpD_Δ29-64_*) was created via overlapping PCR. *E. coli* chromosomal *nlpD* served as template for the PCR, with JR1/JR2 as external primers and JR7/JR8 as internal primers. The truncated allele was then cloned into pAM238 at the same restriction sites as the full-length allele, producing pJR10.

*rcsFFkpA* and *rcsFcol* were obtained by inserting DNA sequences corresponding to helical linker fragments (FkpA Ser94-Glu125 and colicin IA Ile213-Lys282) into *rcsF_Δ1947_* at NcoI and RsrII restriction sites. The *fkpA* gene fragment was amplified from the *E. coli* MC4100 chromosome (JS50/JS51 primers) and the *cia* gene fragment was chemically synthetized as a gene block by Integrated DNA Technologies (IDT). The resulting plasmids were pJS18 and pJS27, respectively. pAM238 does not contain the *lacIq* repressor. Therefore, to enable expression-level regulation by IPTG, strains containing the pAM238 plasmids expressing RcsF variants were co-transformed with pET22b, a high-copy plasmid from a different incompatibility group (pBR223 origin of replication; Novagen) containing the *lacIq* repressor. Chromosomal insertion of *RcsF_Δ19-47_* was performed via λ-Red recombineering^51^ with pSIM5-Tet plasmid (a gift of D. Hughes). In the first step, the cat-sacB cassette was introduced and later replaced by mutant *rcsF*.

The chromosomal *lolCDE* operon was amplified via PCR using primers JS277 and JS278 (adding a C-terminal His-tag to LolE) and then inserted into pBAD33 using the restriction sites PstI and XbaI, resulting in pJR203. The expression level of LolE-His was verified via immunoblotting.

The sequence encoding *lolB* without its N-terminal cysteine was first amplified from the chromosome via PCR using primers JR50/PL387 (adding a C-terminal Strep-tag). It was then cloned into pET28a using the restriction sites XbaI and PstI. *lolA* was amplified using chromosomal *lolA* as PCR template for primers JR30/JR31 (JR31 contains the sequence of a His-tag) and then cloned into pBAD18 using KpnI and XbaI, resulting in pJR48.

The genes encoding Lgt and Lnt were amplified from the chromosome with PCR primers AG389/AG403 and AG393/JR74, respectively. AG403 and JR74 also encode a Myc-tag. PCR products were cloned into pAM238 using KpnI and PstI. Expression levels were verified via immunoblotting (data not shown). *lspA* was amplified with PCR primers JR77/JR78. The PCR product was cloned into pSC213, a modified pAM238 with a ribosome binding site and a C-terminal Flag tag, using NcoI and BamHI. Expression of LspA-Flag was induced by adding 25 μM IPTG. Expression levels were verified with immunoblots (data not shown).

### Cell fractionation and sucrose density gradients

Cell fractionation was performed as described previously^52^ with some modifications. Four hundred milliliters of cells were grown until the optical density at 600 nm (OD_600_) of the culture reached 0.7. Cells were harvested via centrifugation at 6,000 x *g* at 4 °C for 15 min, washed with TE buffer (50 mM Tris-HCl pH 8, 1 mM EDTA), and resuspended in 20 mL of the same buffer. The washing step was skipped with the *Dlpp* strains to prevent the loss of outer membrane vesicles. DNase I (1 mg; Roche), 1 mg RNase A (Thermo Scientific), and a half tablet of a protease inhibitor cocktail (cOmplete EDTA-free Protease Inhibitor Cocktail tablets; Roche) were added to cell suspensions, and cells were passed through a French pressure cell at 1,500 psi. After adding MgCl_2_ to a final concentration of 2 mM, the lysate was centrifuged at 5,000 x *g* at 4 °C for 15 min in order to remove cell debris. Then, 16 mL of supernatant were placed on top of a two-step sucrose gradient (2.3 mL of 2.02 M sucrose in 10 mM HEPES pH 7.5 and 6.6 mL of 0.77 M sucrose in 10 mM HEPES pH 7.5). The samples were centrifuged at 180,000 x *g* for 3 h at 4 °C in a 55.2 Ti Beckman rotor. After centrifugation, the soluble fraction and the membrane fraction were collected. The membrane fraction was diluted four times with 10 mM HEPES pH 7.5. To separate the inner and the outer membranes, 7 mL of the diluted membrane fraction were loaded on top of a second sucrose gradient (10.5 mL of 2.02 M sucrose, 12.5 mL of 1.44 M sucrose, 7 mL of 0.77 M sucrose, always in 10 mM HEPES pH 7.5). The samples were then centrifuged at 112,000 x *g* for 16 h at 10 °C in a SW 28 Beckman rotor. Approximately 30 fractions of 1.5 mL were collected and odd-numbered fractions were subjected to SDS-PAGE, transferred onto a nitrocellulose membrane, and probed with specific antibodies. Graphs were created in GraphPad Prism 9 via spline analysis of the curves representing a mean of three independent experiments.

### Immunoblotting

Protein samples were separated via 10% or 4-12% SDS-PAGE (Life Technologies) and transferred onto nitrocellulose membranes (GE Healthcare Life Sciences). The membranes were blocked with 5% skim milk in 50 mM Tris-HCl pH 7.6, 0.15 M NaCl, and 0.1% Tween 20 (TBS-T). TBS-T was used in all subsequent immunoblotting steps. The primary antibodies were diluted 5,000 to 20,000 times in 1% skim milk in TBS-T and incubated with the membrane for 1 h at room temperature. The anti-RcsF, anti-DsbD, anti-Lpp, anti-NlpD, anti-LolA, and anti-LolB antisera were generated by our lab. Anti-Pal was a gift from R. Lloubès, and anti-His is a peroxidase-conjugated antibody (Qiagen). The membranes were incubated for 1 h at room temperature with horseradish peroxidase-conjugated goat anti-rabbit IgG (Sigma) at a 1:10,000 dilution. Labelled proteins were detected via enhanced chemiluminescence (Pierce ECL Western Blotting Substrate, Thermo Scientific) and visualized using X-ray film (Fuji) or a camera (Image Quant LAS 4000 and Vilber Fusion solo S). In order to quantify proteins levels, band intensities were measured using ImageJ version 1.46r (National Institutes of Health).

### β-galactosidase assay

β-galactosidase activity was measured as described previously^53^. Graphs representing a mean of six experiments with standard deviation were prepared in GraphPad Prism. Expression-level estimations were performed as follows. Cultures used for β-galactosidase activity (0.5 mL per culture) were precipitated with 10% trichloroacetic acid, washed with ice-cold acetone, and resuspended in 0.2 mL Laemmli SDS sample buffer. Samples (5 μL) were subjected to SDS-PAGE and immunoblotted with anti-RcsF antibody.

### SDS-EDTA sensitivity assay

Cells were grown in LB at 37 °C until they reached an OD_600_ of 0.7. Tenfold serial dilutions were made in LB and plated on LB agar supplemented with spectinomycin (100 μg/mL) when necessary. Plates were incubated at 37 °C. To evaluate the sensitivity of the *pal* mutant, plates were supplemented with 0.01% SDS and 0.5 mM EDTA.

### Microscopy image acquisition

Cells were grown in LB at 37 °C until OD_600_ = 0.5. Cells growing in exponential phase were spotted onto a 1% agarose phosphate-buffered saline pad for imaging. Cells were imaged on a Nikon Eclipse Ti2-E inverted fluorescence microscope with a CFI Plan Apochromat DM Lambda 100X Oil, N.A. 1.45, W.D. 0.13 mm objective. Images were collected on a Prime 95B 25 mm camera (Photometrics). We used a Cy5-4050C (32 mm) filter cube (Nikon). Image acquisition was performed with NIS-Element Advance Research version 4.5.

### Protein purification

JR90 cells were grown in LB supplemented with kanamycin (50 μg/mL) at 37 °C. When the culture OD_600_ = 0.5, the expression of cytoplasmic LolB-Strep was induced with 1 mM IPTG. Cells (1 L) were pelleted when they reached OD_600_ = 3 and resuspended in 25 mL of buffer A (200 mM NaCl and 50 mM NaPi, pH 8) containing one tablet of cOmplete EDTA-free Protease Inhibitor Cocktail (Roche). Cells were lysed via two passages through a French pressure cell at 1,500 psi. The lysate was centrifuged at 30,000 x *g* for 40 min at 4 °C in a JA 20 rotor and the supernatant was mixed with Strep-Tactin resin (IBA Lifesciences) previously equilibrated with buffer A. After washing the resin with 10 column volumes of buffer A, LolB-Strep was eluted with 5 column volumes of buffer A supplemented with 5 mM desthiobiotin. LolB-Strep was finally desalted using a PD10 column (GE Healthcare).

Soluble LolA-RcsF_WT_ and LoA-RcsF_*Δ19-47*_ complexes were purified via affinity chromatography as follows. Cells co-expressing LolA either with wild-type RcsF (JR47) or RcsF_Δ19-47_ (JR44) were grown in LB at 37 °C supplemented with 200 μg/mL ampicillin until OD_600_ = 0.5. Protein expression was then induced with 0.2% arabinose. Cells (1 L) were pelleted at OD_600_ = 3 and resuspended in 25 mL of buffer A containing one tablet of protease inhibitor cocktail. Cells were lysed via two passages through a French pressure cell at 1,500 psi. The lysate was centrifuged at 45,000 x *g* for 30 min at 4 °C using a 55.2 Ti Beckman rotor. To obtain the soluble fraction, the supernatant was centrifuged at 180,000 x *g* for 1 h at 4 °C using the same rotor. The supernatant was added to a His Trap HP column (Merck) previously equilibrated with buffer A. The column was washed with 10 column volumes of buffer A supplemented with 20 mM imidazole and LolA-His was eluted using a gradient of imidazole (from 20 mM to 300 mM). The fractions obtained were analyzed via SDS-PAGE; LolA was detected around 25 kDa (data not shown). RcsF variants were detected via immunoblotting with an anti-RcsF antibody. Fractions containing LolA-RcsF variants were pooled, concentrated to 1 mL using a Vivaspin 4 Turbo concentrator (Cut-off 5 kDa; Sartorius), and purified via size-exclusion chromatography with a Superdex S75-10/300 column (GE Healthcare).

### Pull down and transfer of RcsF variants from LolA to LolB

LolB-Strep was incubated at 30 °C for 20 min under agitation with LolA-RcsF_WT_ or with LolA-RcsF_Δ19-47_ (LolA-RcsF_WT_ and LolA-RcsF_Δ19-47_ complexes were purified as described above). The mixture was added to magnetic Strep beads (MagStrep type 3 beads, IBA Life science) previously equilibrated with buffer A and incubated for 30 min at 4 °C on a roller. After washing the beads with the same buffer, LolB-Strep was eluted with buffer A supplemented with 50 mM biotin. Samples were analyzed via SDS-PAGE and LolA and LolB were detected with Coomassie Brilliant Blue (Bio-Rad). RcsF was detected via immunoblotting with an anti-RcsF antibody.

### Structural analysis of lipoproteins

When X-ray, cryo-EM, or NMR structures were available, the missing residues were completed through comparative modeling using MODELLER version 9.22^54^. If no structure of the lipoprotein was available, then the most pertinent analogous structure from proteins belonging to the same PFAM group was used as template for comparative modeling. The linker was defined as the unstructured fragment from the N-terminal Cys of the mature form until the first residue with well-defined secondary structure (α-helix or β-strand) belonging to a globular domain. Short, intermediate, and long linkers had lengths of <12, 12-22, and >22 residues, respectively. Images were generated using UCSF Chimera version 1.13.1^55^.

## LEGENDS FOR FIGURES IN THE EXTENDED DATA

**Extended Data Figure 1.**
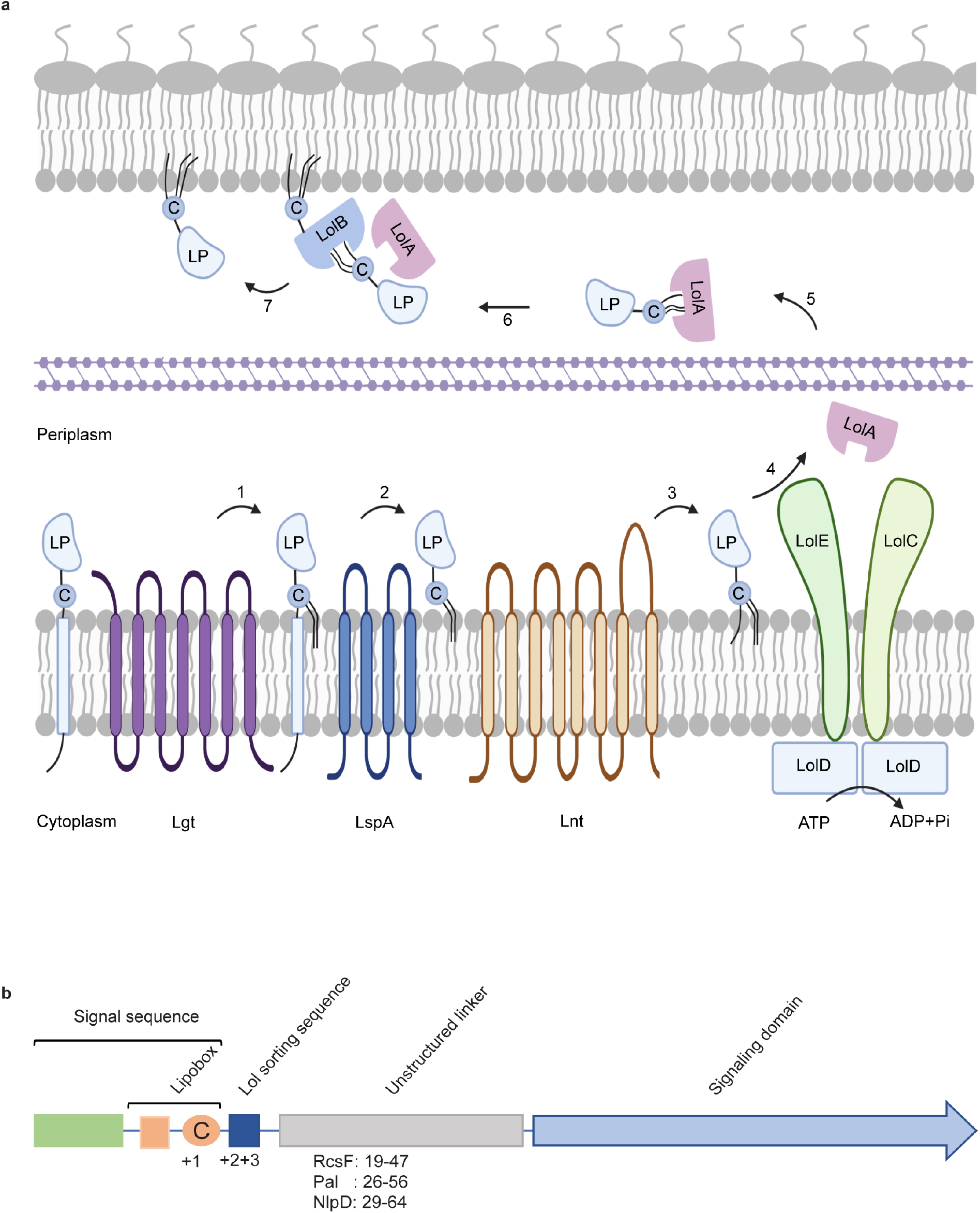
Lipoprotein maturation and sorting in the *E. coli* cell envelope. **a.** After processing by Lgt (step 1), LspA (step 2), and Lnt (step 3), a new lipoprotein either remains in the inner membrane or is extracted by the LolCDE complex (step 4), depending on the residues at position +2 and +3. LolCDE transfers the lipoprotein to the periplasmic chaperone LolA (step 5), which delivers the lipoprotein to LolB (step 6). LolB, a lipoprotein itself, inserts the lipoprotein in the outer membrane using a poorly understood mechanism (step 7). **b.** Schematic of lipoprotein structural domains. The N-terminal signal sequence targets the lipoprotein to the cell envelope; the last four amino acid residues of the signal sequence form the lipobox. The last residue of the lipobox is the invariant cysteine that undergoes lipidation. This cysteine, which is the first residue of the mature lipoprotein, is directly followed by the sorting signal, a sequence of 2 or 3 amino acids that controls the sorting of mature lipoproteins between the inner and outer membranes. The C-terminal portion of a mature lipoprotein is a globular domain. An intrinsically disordered linker separates the sorting signal from the globular domain in about half of *E. coli* lipoproteins (**Fig. 1**; **Extended Data Fig. 2; Extended Data Table 1**). The lengths of the deleted disordered linkers of the unrelated lipoproteins RcsF, Pal, and NlpD are indicated. LP, lipoprotein.

**Extended Data Figure 2.**
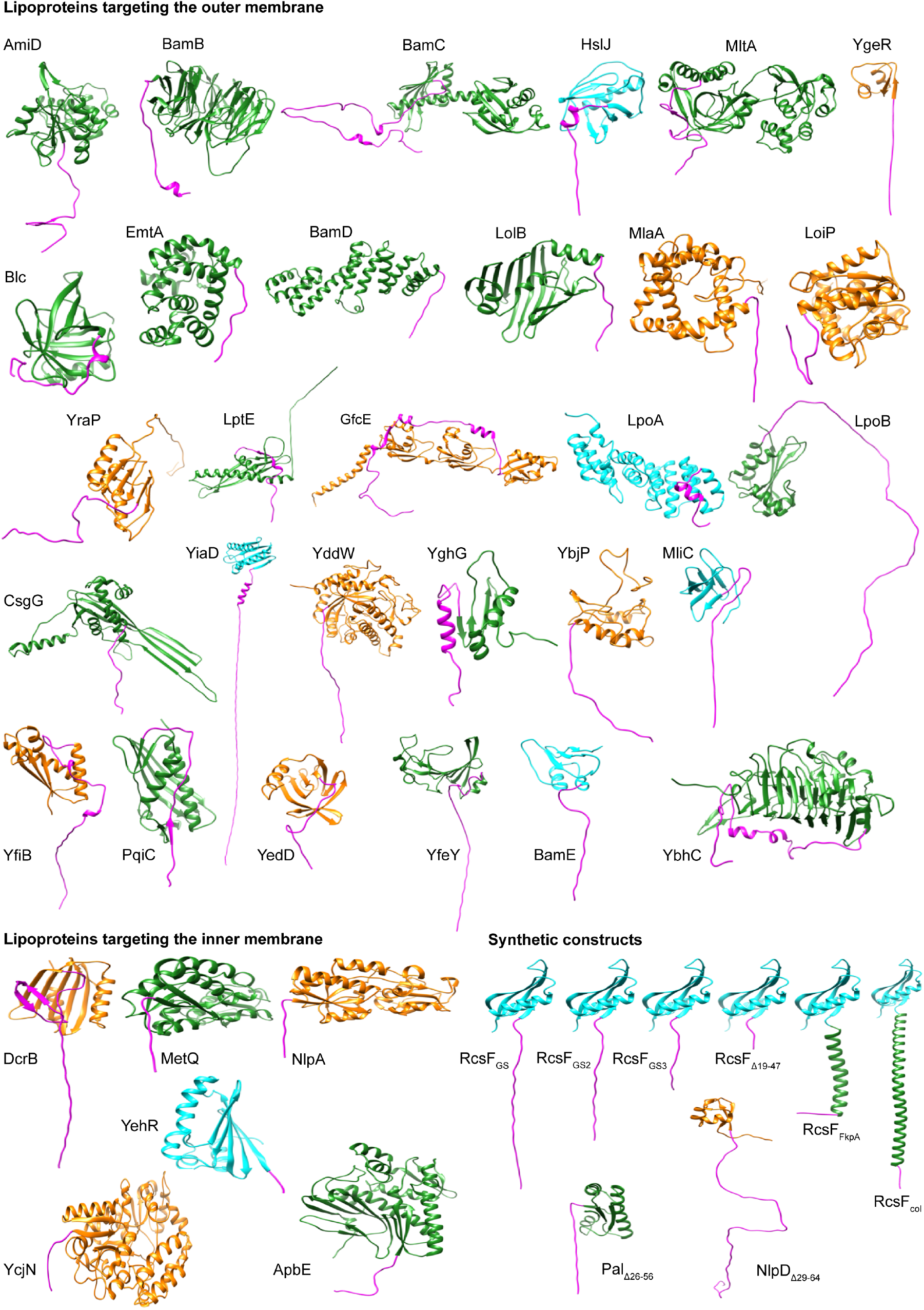
Structural analysis of lipoproteins reveals that half of outer membrane lipoproteins display an intrinsically disordered linker at the N-terminus. Structures were generated via comparative modeling. X-ray and cryo-EM structures are green, NMR structures are cyan, and structures built via comparative modeling from the closest analog in the same PFAM group are orange. In all cases, the N-terminal linker is magenta. Lipoproteins targeting the outer membrane: AmiD, BamB, BamC, HslJ, MltA, LoiP, LpoB, Blc, BamE, CsgG, EmtA, GfcE, BamD, LpoA, LolB, LptE, MlaA, MliC, YddW, YedD, YghG, YfeY, YbjP, YiaD, YbhC, PqiC, YgeR, YfiB, YraP. Lipoproteins targeting the IM: DcrB, MetQ, NlpA, YcjN, YehR, ApbE. Synthetic constructs: RcsF_GS_, RcsF_GS2_, RcsF_GS3_, RcsF_Δ19-47_, RcsF_FkpA_, RcsF_col_, NlpD_Δ29-64_, Pal_Δ26-56_.

**Extended Data Figure 3.**
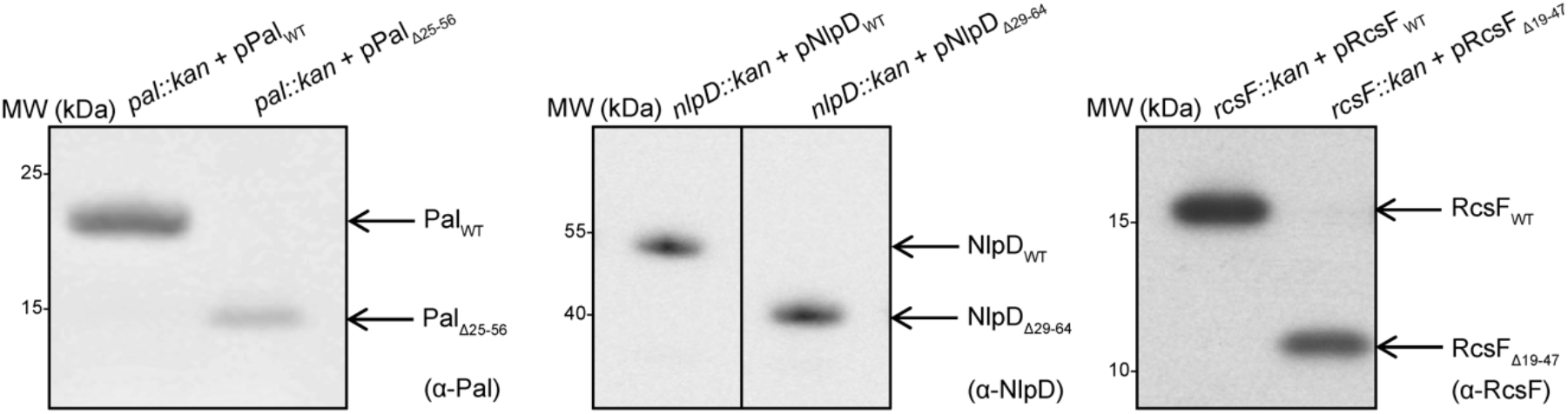
Expression levels of RcsF_Δ19-47_, Pal_Δ26-56_, and NlpD_Δ29-64_. Cells were grown at 37 °C in LB until OD_600_ = 0.5 and precipitated with trichloroacetic acid (Methods). Immunoblots were performed with a-RcsF, a-NlpD, and a-Pal antibodies (Methods). All images are representative of three independent experiments.

**Extended Data Figure 4.**
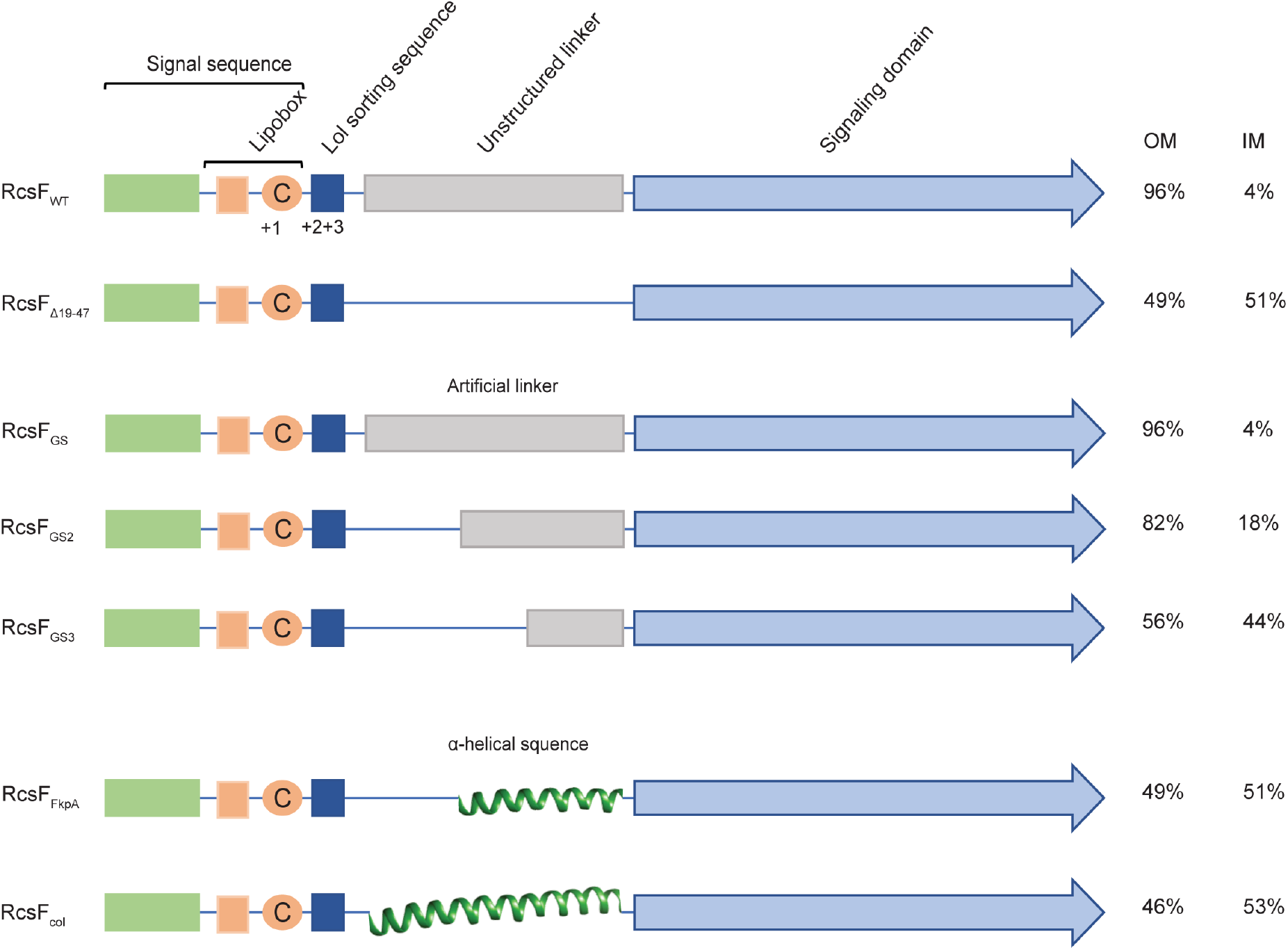
Schematic of RcsF variants used in this study and their distributions in the outer membrane (OM) and inner membrane (IM). RcsF_GS_, RcsF_GS2_, and RcsF_GS3_ have linkers that are disordered and mostly consist of GS repeats. The linker of RcsF_GS_ is the same length as the linker of RcsF_WT_. RcsF_GS2_ and RcsF_GS3_ are shorter than RcsF_WT_. Regions of RcsF_FkpA_ and RcsF_col_ fold into alpha helices borrowed from the sequences of FkpA and colicin Ia, respectively.

**Extended Data Figure 5.**
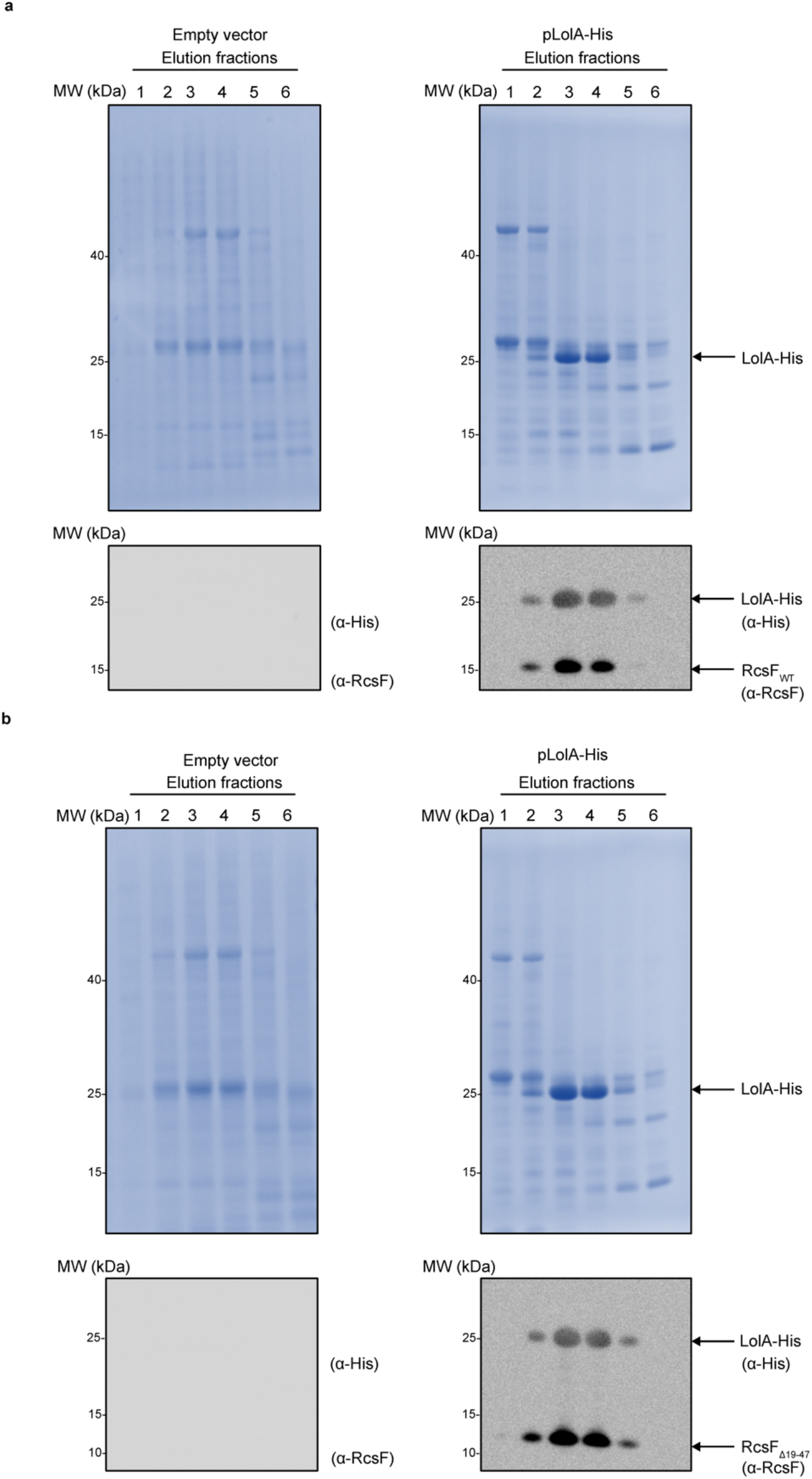
Complexes between LolA and RcsFWT or RcsFΔ19-47 can be purified. Both RcsF_WT_ (a) and RcsF_Δ19-47_ (b) were eluted in complex with LolA-His via affinity chromatography followed by size exclusion chromatography. Gel filtration was performed with a Superdex S75-10/300 column. Samples were analyzed via SDS-PAGE and proteins, including LolA-His, were stained with Coomassie Brilliant Blue (Methods). RcsF variants were detected by immunoblotting fractions with a-RcsF antibodies. Images are representative of three independent experiments.

**Extended Data Figure 6.**
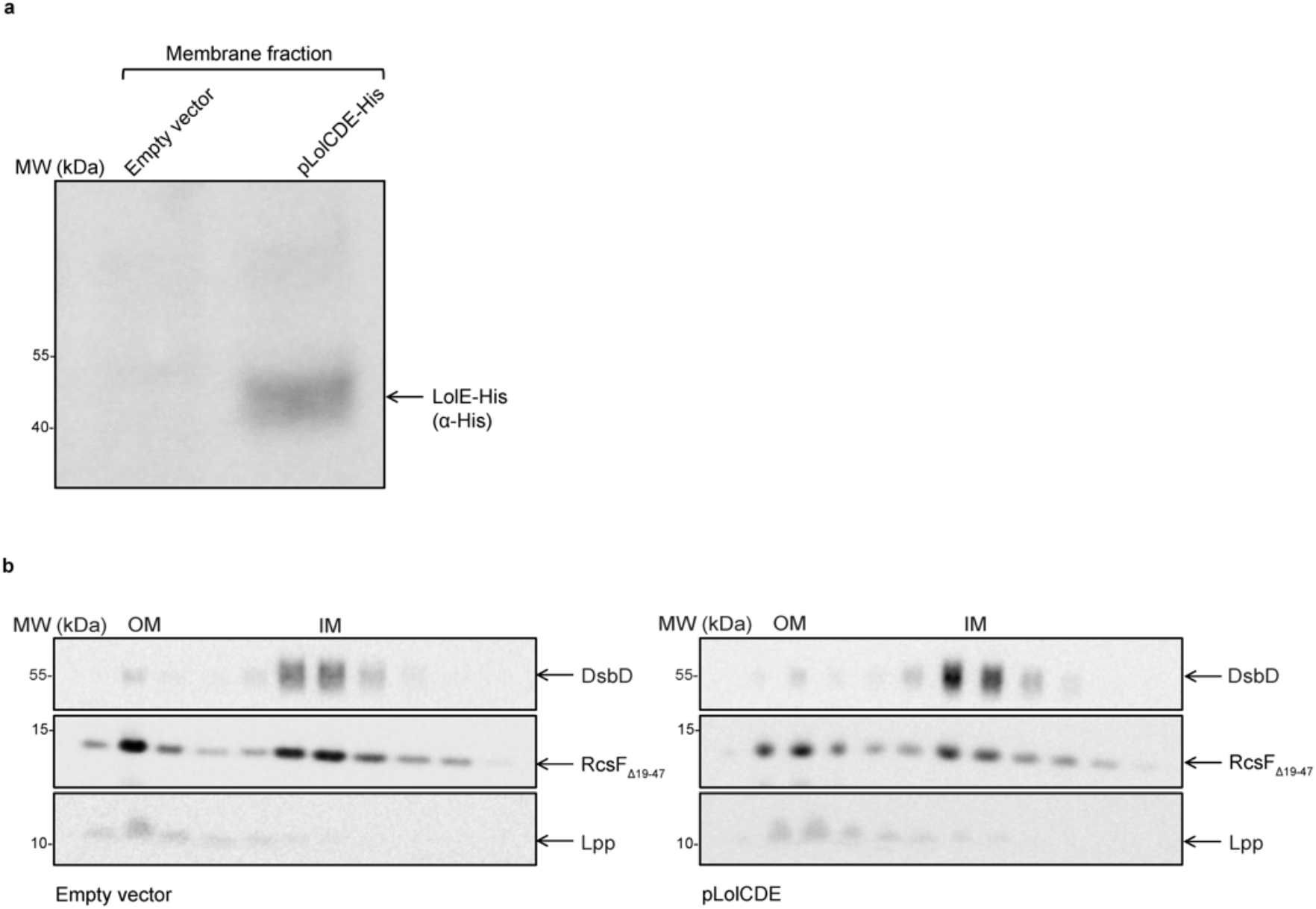
Overexpression of Lol CDE does not restore targeting of RcsF_Δ19-47_. **a.** Expression level of LolCDE-His. Cells were grown in LB plus 0.2% arabinose at 37 °C until OD_600_ = 0.7 (Methods). Membrane and soluble fractions were separated with a sucrose density gradient (Methods). LolE-His was detected in the membrane fraction by immunoblotting with a-His (Methods). Images are representative of three independent experiments. **b.** The outer membrane (OM) and inner membrane (IM) were separated with a sucrose density gradient. Expression of LolCDE did not rescue OM targeting of RcsF_Δ19-47_. Images are representative of experiments performed in biological triplicate.

**Extended Data Figure 7.**
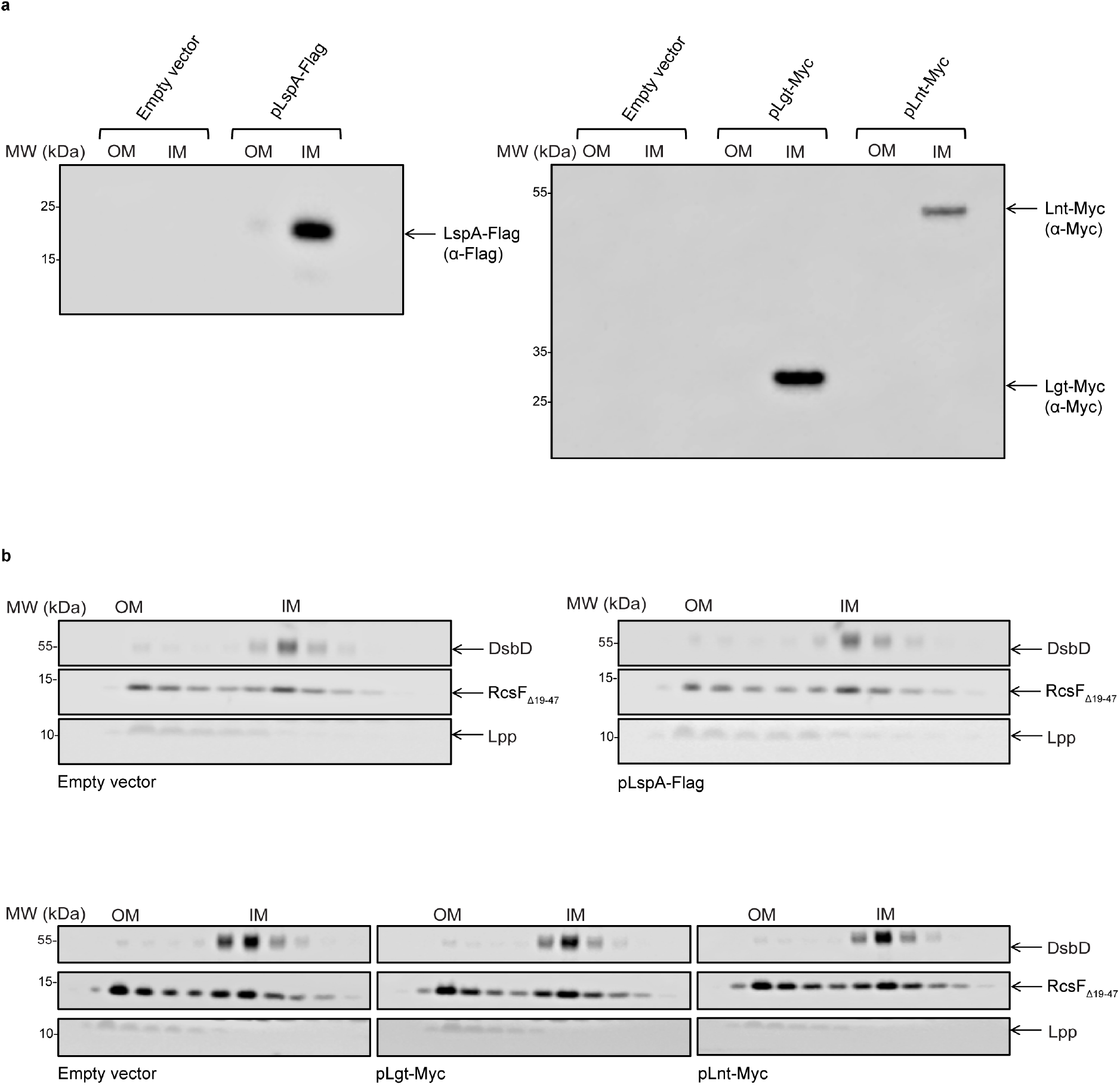
Overexpressing Lgt, LspA, and Lnt does not rescue the targeting of RcsFΔ19-47 to the outer membrane. **a.** Expression levels of Lgt, LspA, and Lnt. Cells were grown in LB (plus 25 μM IPTG for cells expressing LspA) at 37 °C until OD_600_ = 0.7 (Methods). Outer membrane (OM) and inner membrane (IM) were separated with a sucrose density gradient (Methods). Lgt-Myc and Lnt-Myc were detected in the IM via immunoblotting with a-Myc. LspA-Flag was detected in the IM with a-Flag. **b.** Cells overexpressing Lgt, LspA, or Lnt were exposed to a sucrose density gradient (Methods). RcsF_Δ19-47_ was retained in the IM in all conditions. Images are representative of three independent experiments.

## EXTENDED DATA TABLES

**Extended Data Table 1: List of the verified lipoproteins of *E. coli* used for the structural analysis in this study.**

**Attached Excel sheet**

**Extended Data Table 2:**
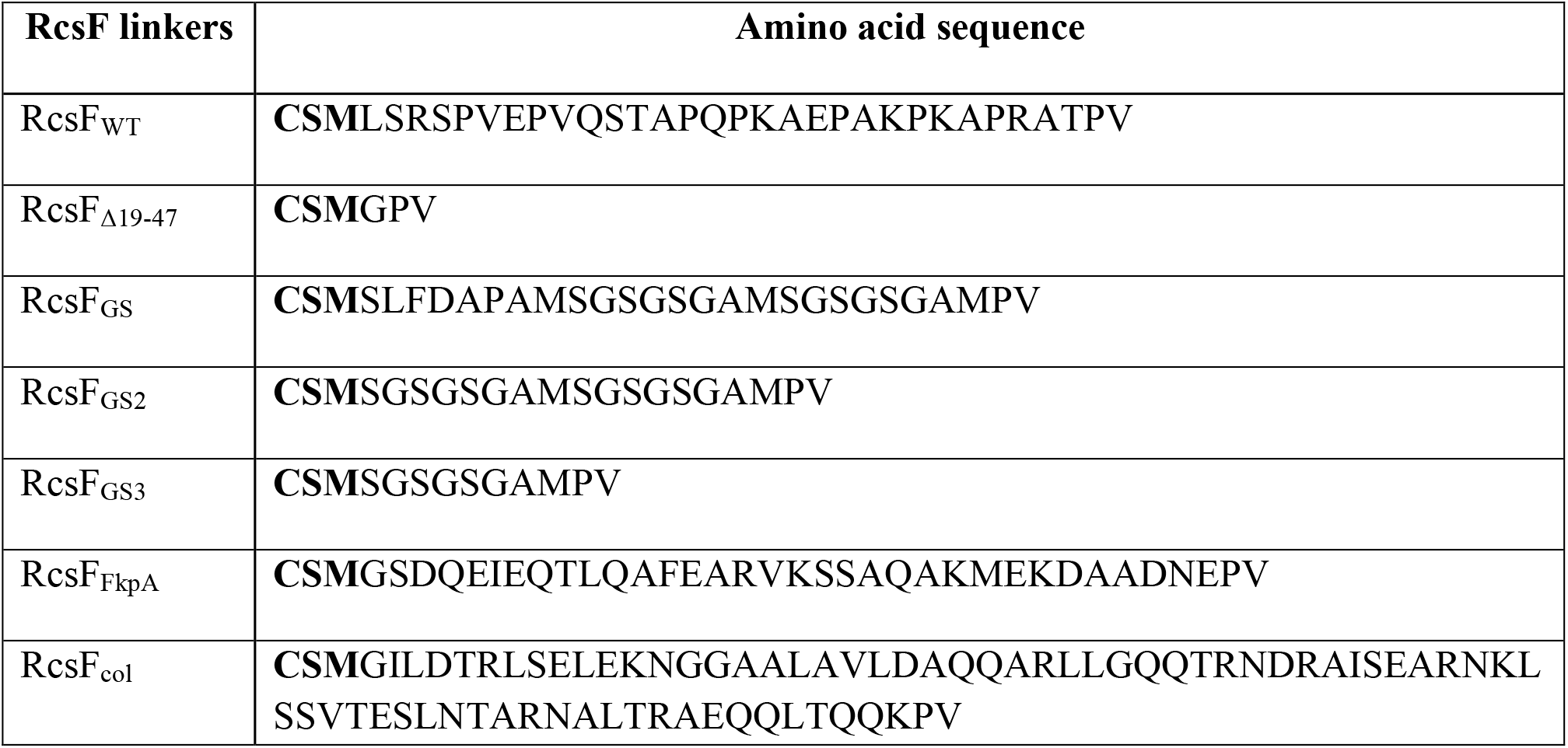
RcsF mutants used in this study and the amino acid sequences of their corresponding N-terminal linkers. The acylated cysteine is the first residue listed.

**Extended Data Table 3:** *E. coli* strains used in this study.

| Strains | Genotype and description | Source |
| --- | --- | --- |
| DH300 | *rprA-lacZ* MG1655 (*argF-lac*) *U169* | <sup>47</sup> |
| Keio collection single mutants | *rcsF::kan, rcsB::kan, pal::kan, nlpD::kan, envC::kan* | <sup>48</sup> |
| XL1-Blue | *endA1 gyrA96 (nal<sup>R</sup>) thi-1 recA1 relA1 lac glnV44F' [::Tn10 proAB<sup>+</sup> lacI<sup>q</sup> Δ(lacZ)M15] hsdR17 (r<sub>K</sub><sup>-</sup> m<sub>K</sub><sup>+</sup>)* | Stratagene |
| BL21 | F- *ompT hsdSB* (rB- mB-) *gal dcm* (DE3) | Novagen |
| JS41 | DH300 Δ*rcsF* pAM238 | This study |
| JS265 | DH300 Δ*rcsF* pJS18 | This study |
| JS346 | DH300 Δ*rcsF rcsB::kan* pET22b | This study |
| JS267 | JS346 pJS18 | This study |
| JS325 | DH300 *pal::kan* | This study |
| JS331 | JS325 pJS20 | This study |
| JS345 | JS325 pJS24 | This study |
| JS360 | DH300 Δ*rcsF* pJS27 | This study |
| JS363 | JS346 pJS27 | This study |
| JS364 | DH300 Δ*rcsF* pSC202 | This study |
| JS372 | DH300 Δ*rcsF* pSC201 | This study |
| JS395 | JS346 pSC198 | This study |
| JS396 | JS346 pSC199 | This study |
| JS397 | JS346 pSC200 | This study |
| JS398 | JS346 pSC201 | This study |
| JS573 | JS346 pSC202 | This study |
| JS574 | DH300 Δ*rcsF* pSC198 | This study |
| JS575 | DH300 Δ*rcsF* pSC199 | This study |
| JS576 | DH300 $\Delta rcsF$ pSC200 | This study |
| JS639 | $\Delta rcsB$ *lpp::kan rcsF::rcsF<sub>Δ19-47</sub>* | This study |
| JR30 | *nlpD::kan* | This study |
| JR31 | JR30 pJR8 | This study |
| JR32 | JR30 pJR10 | This study |
| JR2 | DH300 pAM238 | This study |
| JR88 | BL21 *rcsF::kan* | This study |
| JR90 | JR88 pET28-cytoplasmic LolB-Strep | This study |
| JR187 | *rcsB::kan rcsF::rcsF<sub>Δ19-47</sub>* | This study |
| JR149 | $\Delta nlpD$ | This study |
| JR121 | $\Delta nlpD$ *envC::kan* | This study |
| JR122 | JR121 pJR8 | This study |
| JR123 | JR121 pJR10 | This study |
| JR188 | JR187 pAM238 | This study |
| JR191 | JR187 pAG833 | This study |
| JR204 | JR187 pJR203 | This study |
| JR194 | JR187 pBAD33 | This study |
| JR211 | JR187 pJR209 | This study |
| JR257 | JR187 pJR239 | This study |
| JR274 | JR149 *lpp::kan* | This study |
| JR279 | JR274 pJR10 | This study |
| JR292 | JS325 pAM238 | This study |
| JR293 | JR187 pSC213 | This study |
| JR44 | *rcsB::kan rcsF::rcsF<sub>Δ19-47</sub>* pJR48 | This study |
| JR47 | *rscB::kan* pJR48 | This study |
| JR77 | *rscB::kan rcsF::rcsF<sub>Δ19-47</sub>* pBAD18 | This study |
| JR78 | *rscB::kan* pBAD18 | This study |

**Extended Data Table 4:**
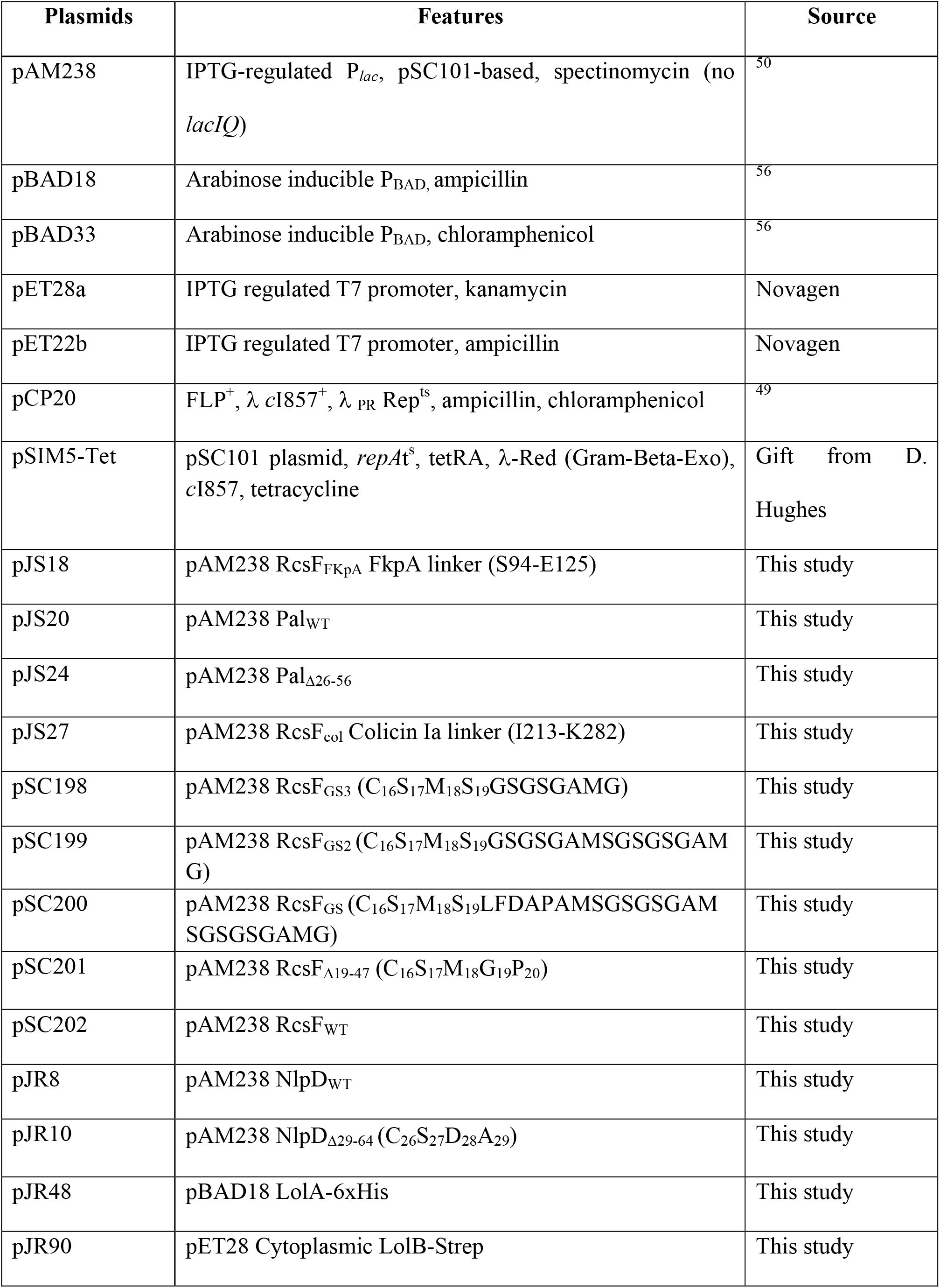
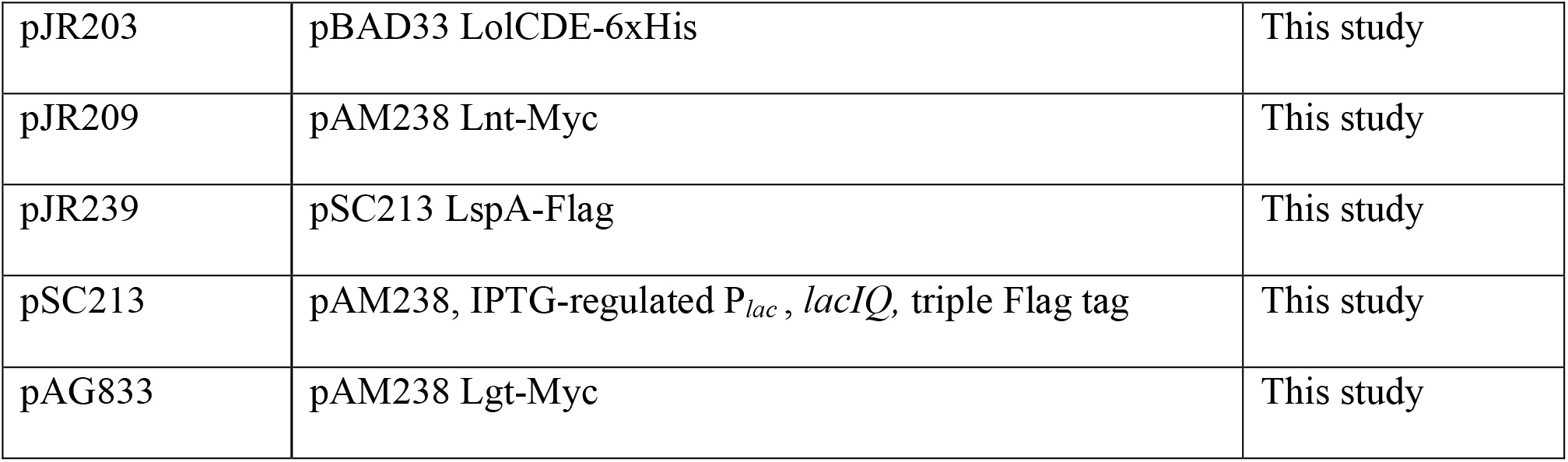
Plasmids used in this study.

**Extended Data Table 5:**
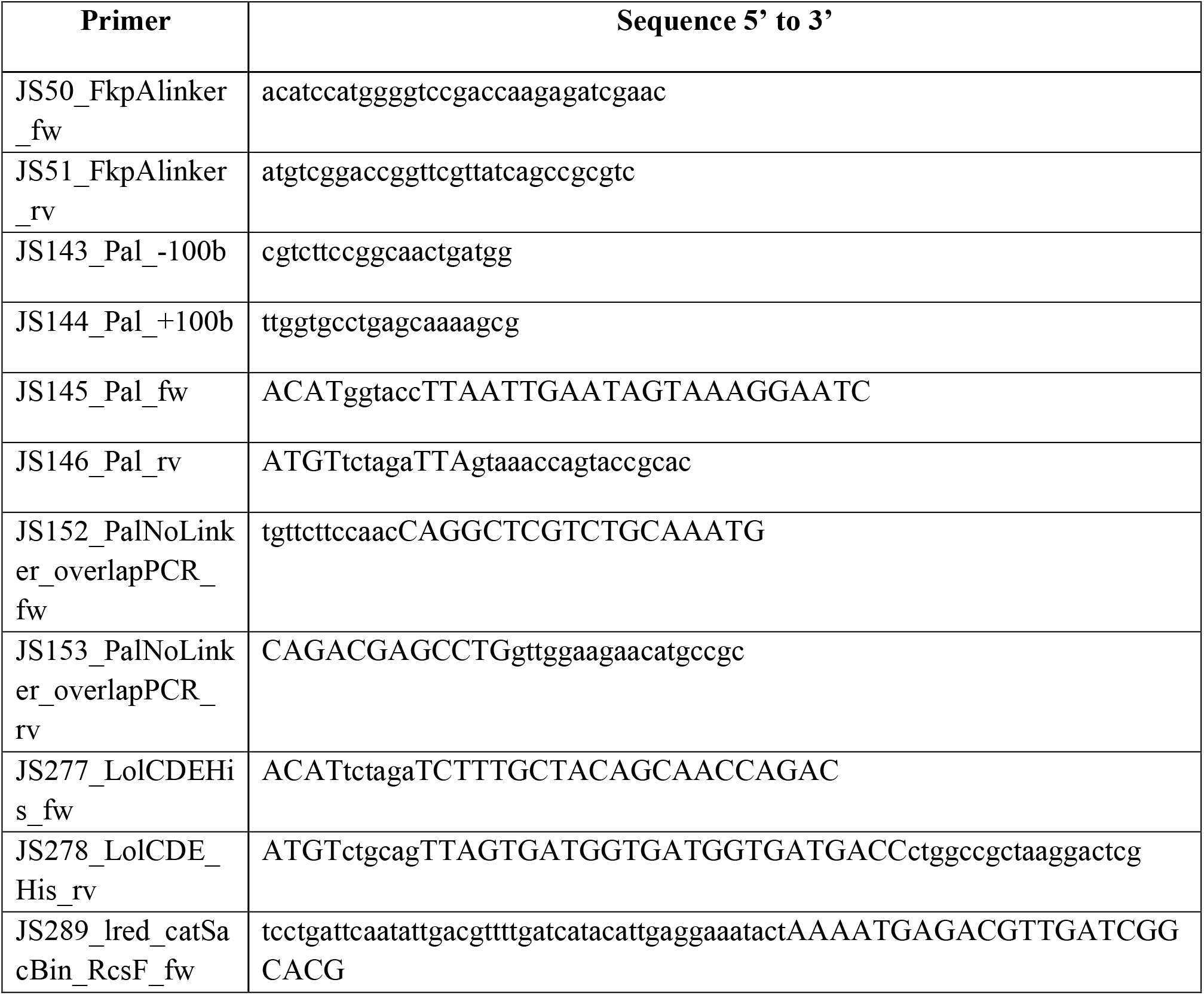
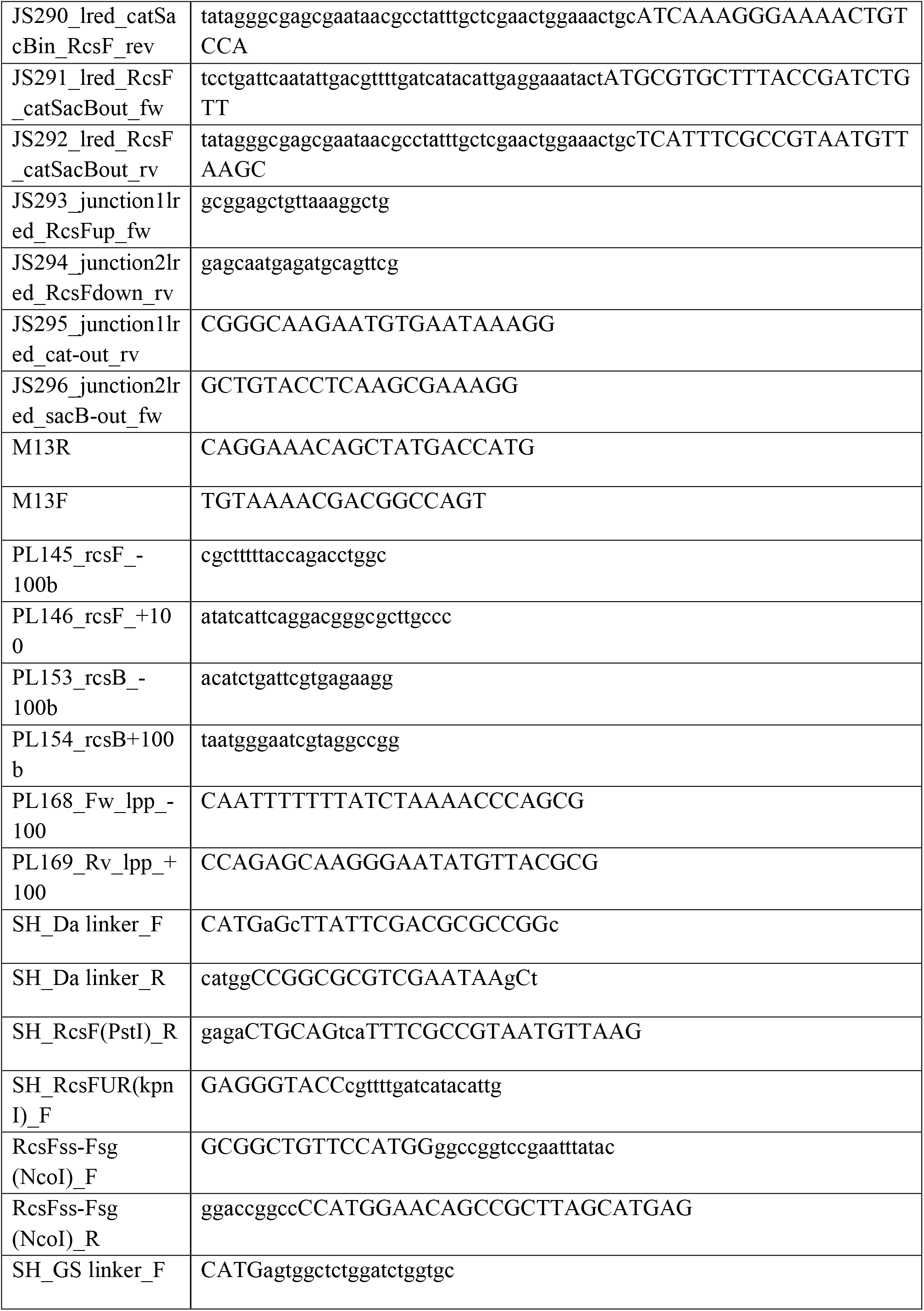
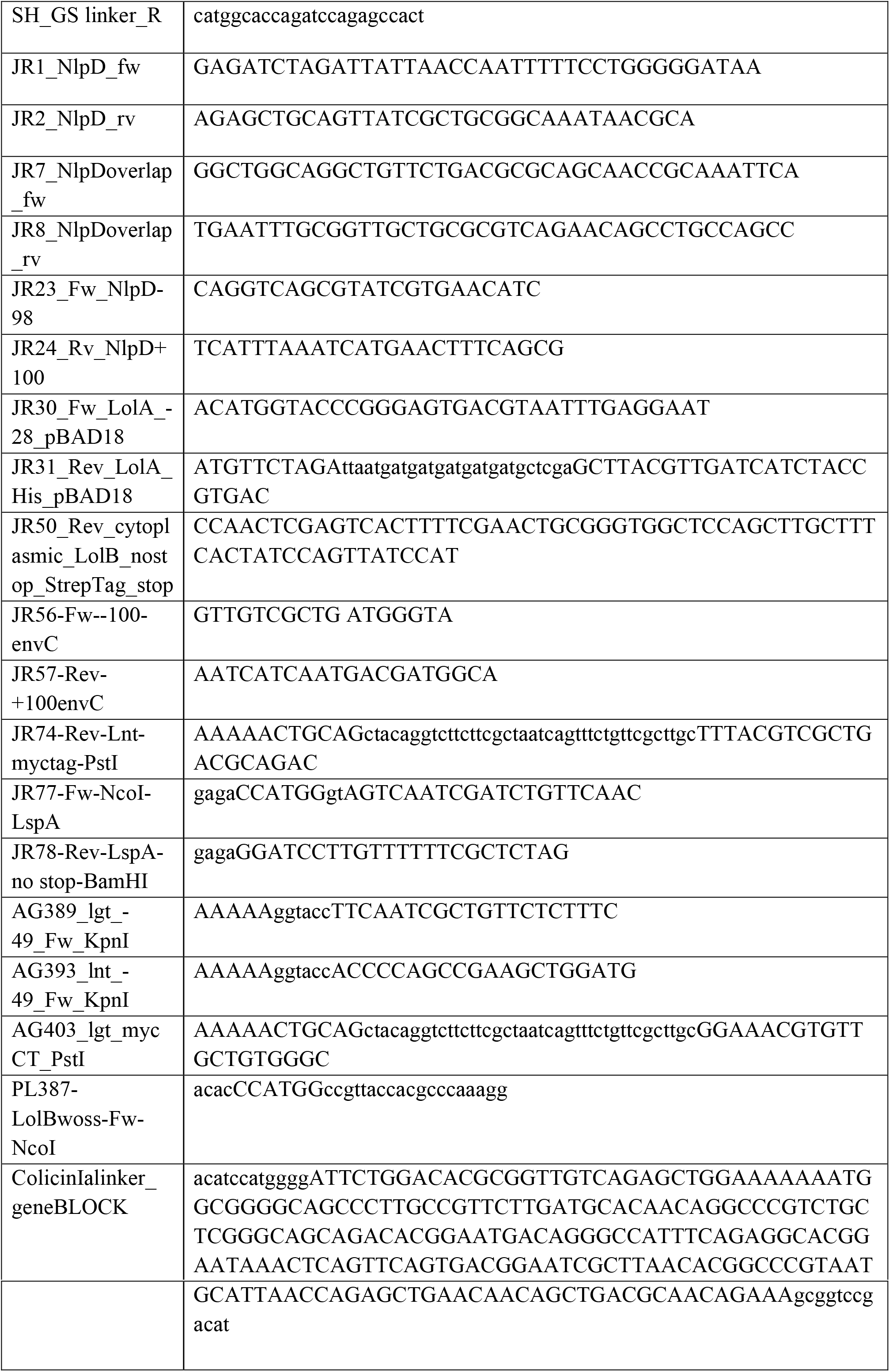
Primers used in this study.

